# Neurotensin receptor allosterism revealed in complex with a biased allosteric modulator

**DOI:** 10.1101/2022.12.26.521971

**Authors:** Brian E. Krumm, Jeffrey F. DiBerto, Reid H. J. Olsen, Hye Jin Kang, Samuel T. Slocum, Shicheng Zhang, Ryan T. Strachan, Lauren M. Slosky, Anthony B. Pinkerton, Lawrence S. Barak, Marc G. Caron, Terry Kenakin, Jonathan F. Fay, Bryan L. Roth

**Affiliations:** Department of Pharmacology, University of North Carolina at Chapel Hill School of Medicine, Chapel Hill, North Carolina, 27599-7365, USA; Department of Pharmacology, University of Minnesota, Minneapolis, Minnesota, 55455, USA; Conrad Prebys Center for Chemical Genomics at Sanford Burnham Prebys Medical Discovery Institute, La Jolla, California 92037, USA; Department of Cell Biology, Duke University, Durham, North Carolina, 27710, USA; Departments of Medicine and Neurobiology, Duke University, Durham, North Carolina, 27710, USA; National Institute of Mental Health Psychoactive Drug Screening Program (NIMH PDSP), School of Medicine, University of North Carolina at Chapel Hill School of Medicine, Chapel Hill, North Carolina 27599-7365, USA; Department of Biochemistry and Biophysics, University of North Carolina at Chapel Hill School of Medicine, Chapel Hill, NC, USA; Division of Chemical Biology and Medicinal Chemistry, Eshelman School of Pharmacy, University of North Carolina at Chapel Hill, Chapel Hill, North Carolina 27599-7360, USA

**Keywords:** GPCR, neurotensin receptor, allosteric, SBI-553

## Abstract

The NTSR1 neurotensin receptor (NTSR1) is a G protein coupled receptor (GPCR) found in the brain and peripheral tissues with neurotensin (NTS) being its endogenous peptide ligand. In the brain, NTS modulates dopamine neuronal activity, induces opioid-independent analgesia, and regulates food intake. Recent studies indicate that biasing NTSR1 toward β-Arrestin signaling can attenuate the actions of psychostimulants and other drugs of abuse. Here we provide the cryoEM structures of NTSR1 ternary complexes with heterotrimeric Gq and Go with and without the brain penetrant small molecule SBI-553. In functional studies, we discovered that SBI-553 displays complex allosteric actions exemplified by negative allosteric modulation for G proteins that are G*α* subunit selective and positive allosteric modulation and agonism for β-Arrestin translocation at NTSR1. Detailed structural analysis of the allosteric binding site illuminated the structural determinants for biased allosteric modulation of SBI-553 on NTSR1. These insights promise to both accelerate the structure-guided design of more effective NTSR1 therapeutics and provide insights into the complexities of GPCR allosteric modulation.

## INTRODUCTION

G protein coupled receptors (GPCRs) comprise the largest family of cell-surface receptors and recognize a diverse array of endogenous and exogenous ligands (Wacker et al, Cell 2017). GPCRs possess a common canonical fold of seven transmembrane (7TM) helical segments connected by three extracellular loops (ECLs) and three intracellular loops (ICLs)(Katritch et al., 2013). Upon agonist binding, GPCRs stabilize conformations that facilitate coupling to heterotrimeric G proteins which in turn elicit second messenger signaling, kinase cascade activation, and ion channel modulation (Pierce et al., 2002). Activated GPCRs can be regulated through mechanisms including phosphorylation by G protein-coupled receptor kinases (GRKs), recruitment of Arrestins, along with desensitization and internalization via clathrin-coated pits. Arrestin binding in turn can facilitate the activation of a number of downstream signaling events thereby altering the duration and magnitude of signal transduction (Lefkowitz and Shenoy, 2005). Thus, Arrestins serve as scaffolds or adapters recruiting a variety of signaling molecules to activated GPCRs connecting them to various intracellular signaling pathways including MAPK cascades while also interacting with and modifying the activity of cytoplasmic transcription factors (Wilbanks et al., 2004).

Allosteric modulators for GPCRs have been known for many decades beginning with the identification of negative allosteric modulators (NAMs) like sodium ion at opioid receptors (Pert et al., 1973) (Simon and Groth, 1975). Later, both positive allosteric modulators (PAMs) and NAMs at muscarinic cholinergic receptors (Clark and Mitchelson, 1976; Kenakin and Boselli, 1989) (Christopoulos and Mitchelson, 1997) were identified. Mechanistically, PAMs augment the affinity and/or efficacy of orthosteric agonists, while NAMs reduce these actions. The structural basis for GPCR positive and negative allosteric modulation has been described for several GPCRs (see (Thal et al., 2018) for review) with the first being structures of GPCRs complexed with the NAM sodium ion (Liu et al., 2012) (Fenalti et al., 2014; Wang et al., 2017). To our knowledge, there are no structures of GPCRs in complex with Arrestin biased allosteric modulators and transducer. Accordingly, the structural basis of allosteric modulation remains incompletely elucidated.

Recently, a series of small molecules for the Neurotensin Receptor 1 (NTSR1) were identified via High Content Screening (HCS) using a β-Arrestin translocation assay (Barak et al., 2016; Peddibhotla et al., 2013) (Hershberger et al., 2014). Members of this series were shown to bias NTSR1 towards β-Arrestin signaling by a mechanism consistent with allosterism. Unfortunately, poor aqueous solubility and unfavorable pharmacokinetic profiles precluded clinical formulation along with the structural and animal studies needed to elucidate their mechanism of action. Subsequent medicinal chemistry optimization yielded SBI- 553, a brain penetrant and bioavailable (in rodents) small molecule compound (Pinkerton, et al., 2019). Recent animal studies suggest SBI-553 is a unique allosteric modulator, exhibiting β-Arrestin agonism on its own while also conferring β-Arrestin bias to the endogenous neurotensin peptide (NTS) of NTSR1 (Slosky et al., 2020). Thus, Slosky, L.M, et al (2020) show that this biasing of NTSR1 functional selectivity translates into directed pharmacological actions *in vivo*, with the attenuation of psychostimulant-associated behaviors devoid of the side effects characteristic of unbiased NTSR1 agonism. Here, through detailed functional characterization, we show that SBI-553 is a novel biased allosteric modulator that stabilizes NTSR1 in a conformationally selective state permissive for activation of only a subset of transducers (β-Arrestin, Gi1, and G12 vs. Gq and G15). To elucidate the structural basis for this unusual mode of allosteric modulation, we determined the structures of NTSR1 in complex with Neurotensin (8-13) (NTS_8-13_) the active portion of the longer endogenous peptide (NTS 1-13), MiniGq (Kim et al., 2020) and MiniGo (Zhang et al., 2022a) (Zhang et al., 2022b) with and without SBI-553 using cryo-electron microscopy (cryoEM) (Figure 1A). As NTSR1 is responsible for mediating neurotensin’s actions in hypothermia, hypotension, antinociception, cancer cell growth, Parkinson’s disease (Bissette et al., 1976) (Schimpff et al., 2001) (Alifano et al., 2010a; Alifano et al., 2010b), and drug abuse (see (Slosky et al., 2020), elucidating the molecular mechanism(s) of action of this unusual allosteric modulator could accelerate basic and translational science.

**Figure 1.**
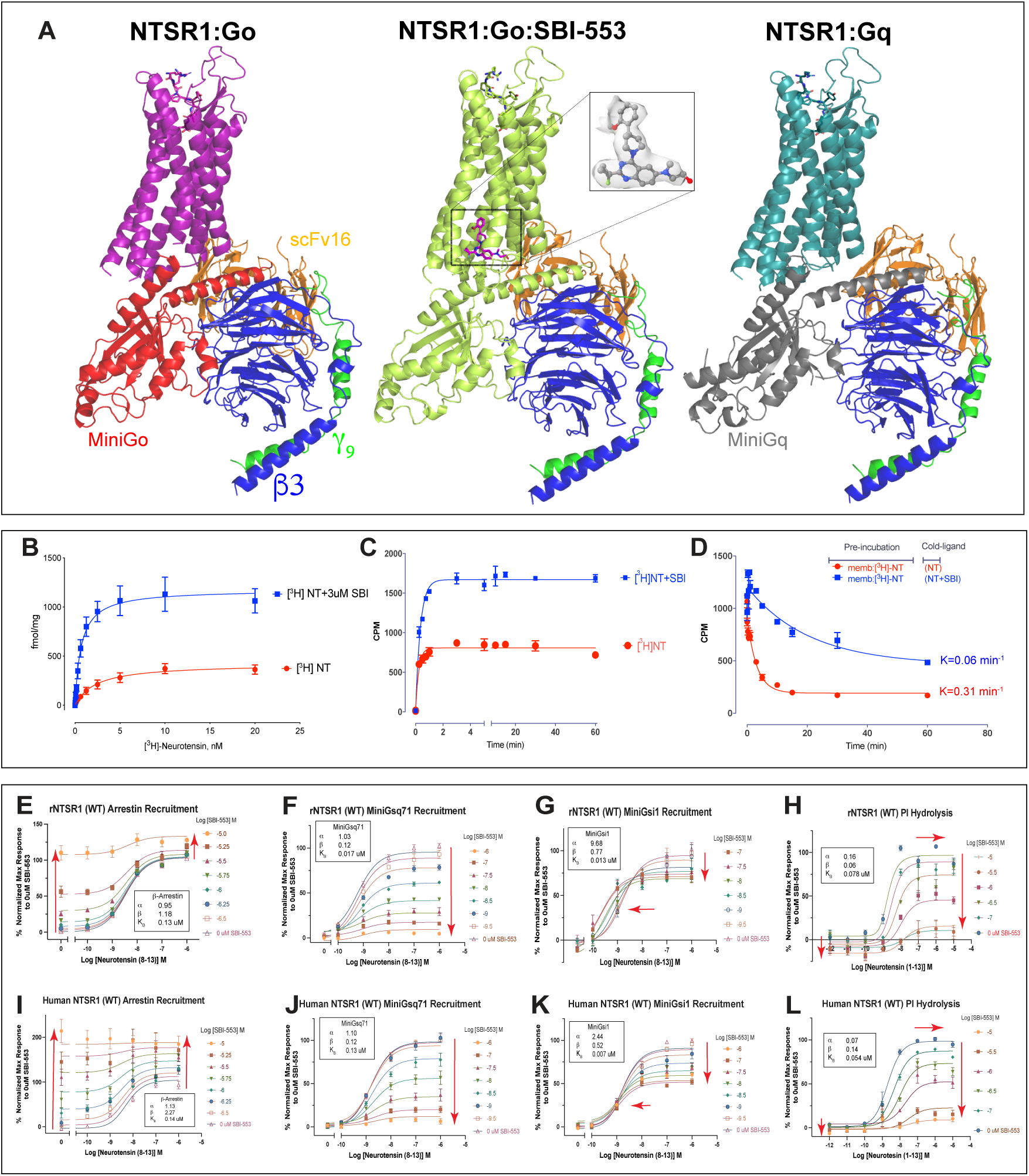
NTSR1 CryoEM Structures and SBI-553 biased allosteric effect on NTSR1 using saturation binding and recruitment assays. **A) Overview of** discussed cryoEM structures of NTSR1 in complex with MiniGo +/- SBI-553 and MiniGq. Insert is representative EM density for SBI-553. In stick figures is SBI- 553 with the allosteric site located in the rNTSR1 intracellular cavity. B) Rat Neurotensin Receptor 1 (rNTSR1) saturation binding experiments with increasing concentrations of [^3^H]Neurotensin (NTS) in the presence (blue) and absence (red) of a constant concentration of 3 μM SBI-553. SBI-553 significantly enhanced [^3^H]NTS binding affinity from 2.3 +/- 0.6 nM to 0.67 +/- 0.14 nM (p=0.036; F(1,92) = 4.53). B) rNTSR1 on-rate and C) rNTSR1 off-rate experiments using 1nM [^3^H]NTS measured at different time intervals in the presence (blue) and absence (red) of constant concentration of 3 uM SBI-553, 10 uM cold NTS was used for off-rate experiments. E) and I) Rat and Human NTSR1 SBI-553 dose response curves for arrestment recruitment using BRET assay. F) and J) SBI-553 dose response curves for rNTSR1 MiniGsq-71 recruitment using BRET assay G) and K) SBI-553 dose response curves for rNTSR1 and hNTSR1 MiniGsi1 recruitment using BRET assay. H) and L) SBI- 553 dose response curves for rNTSR1 and hNTSR1 PI hydrolysis assay. Insert in each graph are the allosteric parameters required to fit the data using the Allosteric EC50 Shift and Black Leff Ehlert equation for PAM, General Least squares fit in Graphpad Prism 8.0 (Graphpad Software Inc., San Diego, CA). Data are presented as mean values ± SEM with a minimum of two technical replicates and N = 3 biological replicates. Red arrows indicate relative effect on fitted dose-response curves with the addition of SBI-553.

## RESULTS

### SBI-553 is a PAM-agonist and biased allosteric modulator

Extensive medicinal chemistry optimization of high content screening (HCS) hits yielded SBI-0654553 (SBI-553), an allosteric modulator with the ability to confer β-Arrestin bias to NTSR1 having improved solubility and efficacy when compared to its parent series (Pinkerton, et al. 2019) (Barak et al., 2016) (Hershberger et al., 2014; Peddibhotla et al., 2013), albeit the mechanism(s) and structural determinants by which SBI-553 modulates NTSR1 signaling remain obscure. Additionally, prior studies postulated, but did not reveal, that these compounds were allosteric modulators with regards to additional NTSR1 signaling pathways.

The mechanisms driving allosteric modulation can be multifactorial, therefore we evaluated SBI-553’s effects using a combination of radioligand binding and functional assays (Langmead, 2011). We investigated the effects of SBI-553 on NTSR1 (note – NTSR1 is understood here to represent both human and rat NTSR1 unless otherwise identified) in radioligand binding assays and found that SBI-553 increased the total number of binding sites (B_max_) of [^3^H]NTS at wild-type rat NTSR1 (rNTSR1) approximately 2.8-fold (Figure 1B). SBI-553 also enhanced [^3^H]NTS binding affinity (K_d(rNTSR1)_ = 2.3 nM, K_d(rNTSR1/SBI)_ = 0.67 nM, K_d(hNTSR1)_ = 2.14 nM, K_d(hNTSR1/SBI)_ = 0.92 nM). In rNTSR1 kinetic experiments, SBI-553 accelerated the association rate and slowed the dissociation rate (Figure 1C and 1D) when added to membranes simultaneously with [^3^H]NTS. These increases in B_max_ and altered kinetic parameters implied an allosteric interaction between NTS and SBI-553.

As SBI-553 was identified based on its ability to induce Arrestin translocation in HCS via NTSR1, we next asked if these effects of SBI-553 on NTSR1 binding kinetics required Arrestin and/or G proteins. For these studies we used cells in which Arrestin or Gq/11 were deleted via CRISPR (Alvarez-Curto et al., 2016). Genetic deletion of *β*Arr1 and *β*Arr2 did not attenuate SBI- 553’s effect to enhance NTS binding for either rat or human NTSR1 (Figure S1A). As NTSR1 is reported to interact with Gq, we also assessed the effect of genetic deletion of Gq and G11 on SBI-553’s enhancement of radioligand binding. No inhibition of SBI-553’s effect was observed indicating that SBI-553 actions are Gq/11-independent (Figure S1A). Taken together, these results imply that the allosteric actions of SBI-553 on radioligand binding are Arrestin and G protein-independent. Finally we note that the effects of SBI-553 are consistent with predictions from the Hall model for allosteric binding (Hall, 2000) which describes how allosteric modulation in the absence of transducers can modulate apparent K_d_ (equilibrium dissociation constant) and B_max_ values for agonists.

We next investigated the mechanism(s) for this potential allosteric modulation in functional assays. Importantly, both the allosteric ternary complex model (ATCM) and the allosteric operational model define the overall allosteric effect of the modulator on orthosteric agonist affinity by the alpha value (α), while the overall allosteric effect of the modulator on the orthosteric agonist efficacy is defined by the beta value (β). Values of α > 1 indicate positive cooperativity, values of 0 < α < 1 indicate diminished cooperativity, and α values = 1 indicate neutral (silent) cooperativity. Additionally, changes in efficacy when β > 1 define a PAM effect, and values of 0 < β < 1 define a NAM effect. Additional allosteric parameters are the equilibrium dissociation constants of the agonist and allosteric modulator–receptor complexes, K_A_ and K_B_, respectively. The receptor and orthosteric agonist complex efficacy is defined by the τ_A_ term, while the receptor and allosteric agonist complex efficacy is defined by the τ_B_ term. In practice, a PAM increases the GPCR’s response to the orthosteric agonist, whereas a NAM inhibits the GPCR’s response to the orthosteric agonist. Unique among these allosteric modulators is the PAM-Antagonist, which targets agonist-activated receptors (α > 1) to shut them down (β < 1) (Kenakin and Strachan, 2018) .

We next performed bioluminescence resonance energy transfer (BRET) experiments to more completely understand SBI-553 modulator effects on NTSR1 *vis-a-vis* agonist-induced *β−*Arrestin and G protein recruitment (Figure S1C for visual depiction of assay). In β−Arrestin recruitment assays (Figures 1E and 1I, Tables S2/S3), SBI-553 has both direct agonist activity in the absence of NTS and PAM activity in conjunction with NTS. Thus, for βArr recruitment SBI- 553 is classified as a PAM-agonist (Christopoulos et al., 2014). Quantification of the effect of SBI-553 on NTS concentration-response curves was performed using the functional allosteric model (Ehlert, 2005) (Price et al., 2005) (Watson et al., 2005) which is a combination of the Stockton-Ehlert allosteric binding model (Stockton et al., 1983) (Ehlert, 1988) and the Black/Leff operational model (Black and Leff, 1983). This analysis revealed that SBI-553 significantly potentiates the efficacy of NTSR1 for recruiting Arrestin in the absence of NTS at concentrations of 3 μM (τ_B(rNTSR1)_ (3 μM) = 0.89, τ_B(hNTSR1)_ (3 μM) = 1.06 (hNTSR1) although this potentiation diminishes expectedly at lower concentrations of SBI-553. In terms of allosteric effects, SBI-553 in a cooperative manner potentiated the efficacy of NTSR1 for NTS albeit while also acting in a mostly cooperative manner with NTS affinity (α = 0.95, β = 1.18, (rNTSR1), α = 1.13, β = 2.27 (hNTSR1), Tables S2/S3). Conversely, in the BRET Gq recruitment assay (Figures 1F, 1J, and 2A), SBI-553 displayed characteristic PAM-Antagonist activity by enhancing NTS affinity while at the same time antagonizing NTS efficacy (α = 1.03, β = 0.12 (rNTSR1), α = 1.10, β = 0.12 (hNTSR1), Tables S2 and S3). Given SBI-553’s complex actions on enhancing agonist binding, slowing dissociation, enhancing agonist activity for Arrestin and negative allosteric modulation of Gq recruitment, we characterize SBI-553 as a biased allosteric modulator with βArr PAM-agonism and Gq-specific PAM-antagonism (Christopoulos et al., 2014) (Kenakin and Strachan, 2018) .

### SBI-553 is a biased allosteric modulator with diverse G protein specific effects

NTSR1 is known to couple to multiple effectors of G protein signaling pathways including Gq, Gi/o and to a much lesser extent Gs (Grisshammer and Hermans, 2001) Therefore, we considered the possibility that SBI-553 could exert more complex allosteric actions on NTS signaling to multiple pathways (Kenakin and Christopoulos, 2013). To test this possibility, we initially confirmed SBI-553’s ability to negatively modulate NTSR1’s activation of Gq using a BRET2 G protein dissociation assay (Figure S1C for graphic depiction of BRET2 assay). In the BRET2 assay, activation of NTSR1 resulted in a concentration-dependent decrease in net BRET signal due to nucleotide exchange and heterotrimer dissociation. Consistent with a negative modulation of Gq protein recruitment, SBI-553 exhibited a pronounced dose-dependent NAM effect on Gq activation (α = 0.07, β = 0.49 (rNTSR1), α = 0.06, β = 0.41 (hNTSR1), Figures 2A, 3A and 3B, and Tables S2/S3).

**Figure 2.**
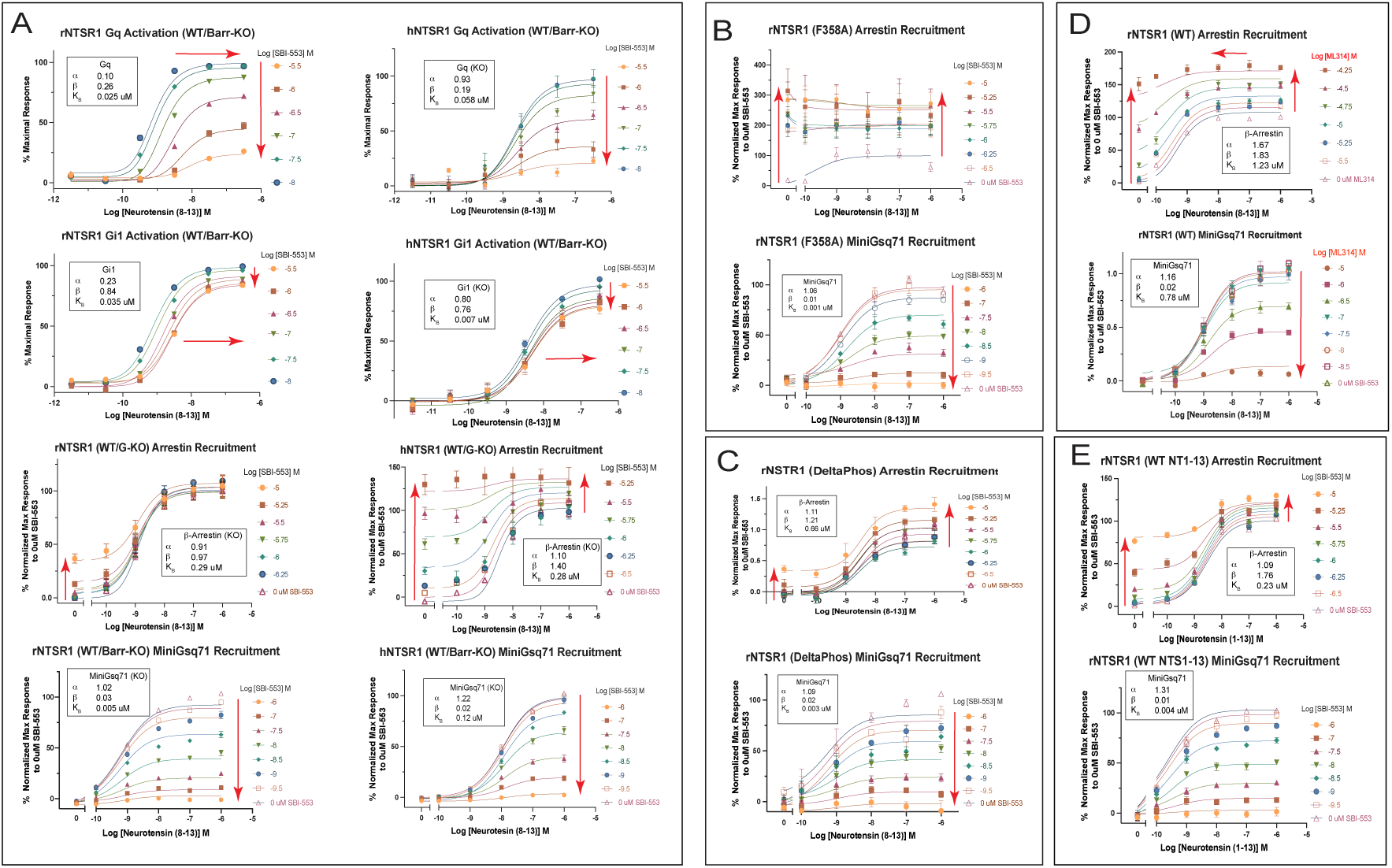
Functional characterization of SBI-553 effects on rNTSR1 and hNTSR1 BRET activation and recruitment assays. **A)** Functional characterization of SBI-553 effects on hNTSR1 using BRET, BRET2 assays in knockout cell lines as described in the main text. B) Effects of NTSR1 constitutive activity on SBI-553 signal modulation. C) Effects of NTSR1 phosphorylation sites deletions on SBI-553’s signal modulation. D) Comparison of SBI-553 parent compound (ML-314) dose response curves on Arrestin and MiniGsq recruitment. E) Comparison of full length NTS (1-13) dose response in the presence of SBI-553 on Arrestin and MiniGsq recruitment. Data are presented as mean values ± SEM with a minimum of two technical replicates and N = 3 biological replicates. Insert in each graph are the allosteric parameters required to fit the data using the Allosteric EC50 Shift and Black Leff Ehlert equation for PAM, General Least squares fit in Graphpad Prism 8.0 (Graphpad Software Inc., San Diego, CA). Red arrows indicate relative effect on fitted dose- response curves with the addition of SBI-553.

**Figure 3.**
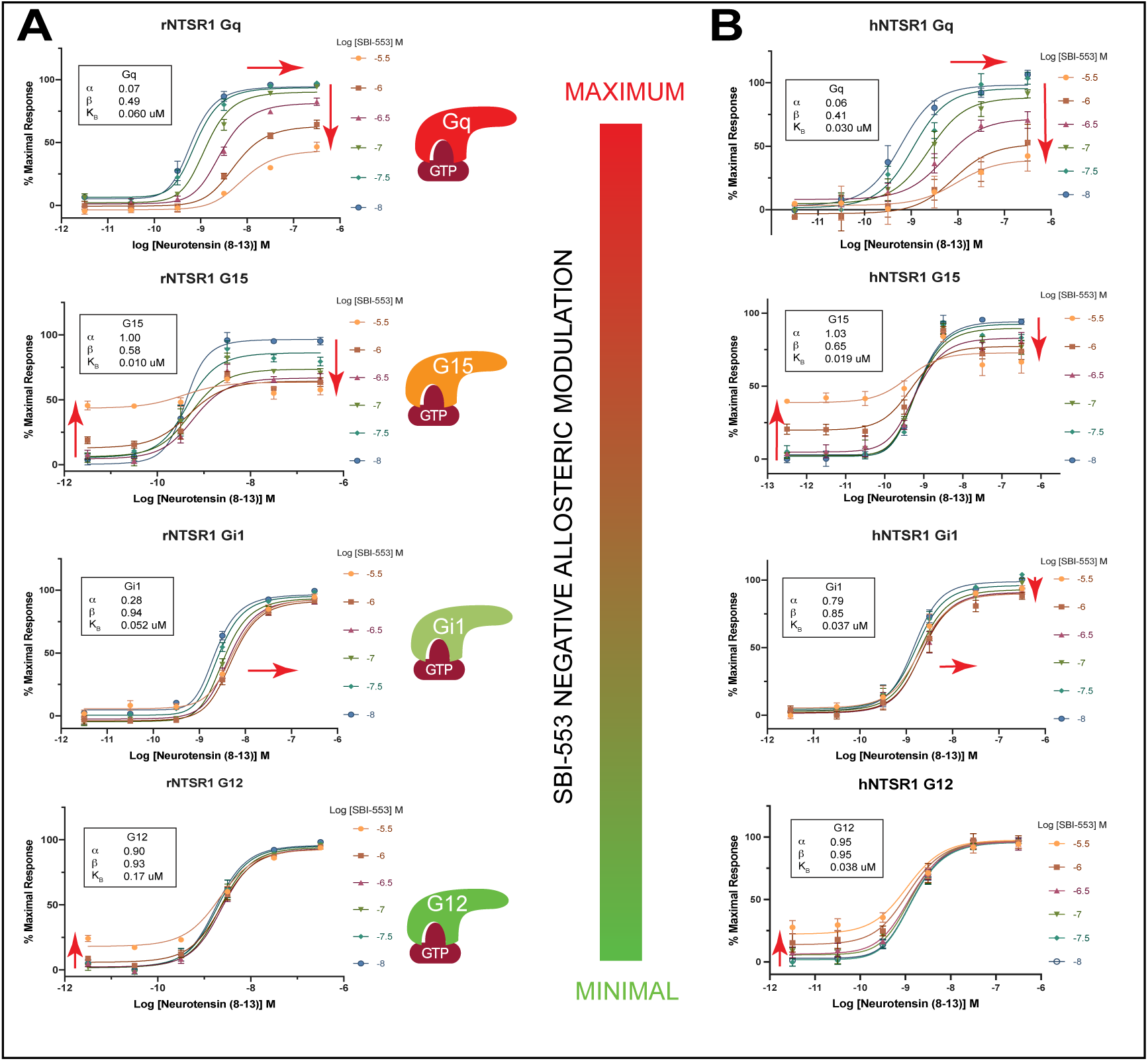
SBI-553 is a biased allosteric modulator with diverse G protein- specific effects. A) rNTSR1 and B) hNTSR1 G protein activation assay (BRET2) SBI-553 dose response curves in the presence of NTS_8-13_ reveal differential effects on signaling with maximum effect on Gq signaling and minimal effect on G12 signaling. Insert in each graph are the allosteric parameters required to fit the data using the Allosteric EC50 Shift and Black Leff Ehlert equation for PAM, General Least squares fit in Graphpad Prism 8.0 (Graphpad Software Inc., San Diego, CA). Data are presented as mean values ± SEM with a minimum of two technical replicates and N = 3 biological replicates. Red arrows indicate relative effect on fitted dose-response curves with the addition of SBI-553.

We next examined SBI-553’s ability to modulate NTS signaling at other signaling pathways - a set of experiments that revealed the most diverse array of biased allosteric effects seen to date. Specifically, SBI-553’s modulation of NTS G15 activation was consistent with a NAM-agonist profile (Figures 2A, 3A and 3B, Tables S2/S3), displaying agonist activity at high concentrations of SBI-553 and low concentration of NTS (τ_B(rNTSR1)_ = 0.30 and τ_B(hNTSR1)_ = 0.50), and a saturable NAM effect on NTS G15 efficacy (β = 0.58 (rNTSR1), β = 0.68 (hNTSR1)). At Gi1, SBI-553 acted as a very weak NAM-agonist as it displayed very little agonist activity at high concentrations of modulator and low concentrations of NTS (τ_B(rNTSR1)_ = 0.04, τ_B(hNTSR1)_ = 0.07) and exerted a small effect on NTS efficacy (β = 0.94 (rNTSR1), β = 0.85 (hNTSR1)). In contrast, SBI- 553 acted primarily as an allosteric agonist having no apparent effect on G12 activation in the presence of high concentrations of NTS (β = 0.93 (rNTSR1), β = 0.95 (hNTSR1)) but had apparent agonist activity at high concentrations and in the absence of NTS (τ_B(rNTSR1)_ = 0.28, τ_B(hNTSR1)_ = 0.30) (Figures 3A and 3B, Tables S2/S3).

Since SBI-553 enhances recruitment of βArr to NTSR1 it was conceivable that negative modulation of some G protein pathways was due to βArr recruitment and diminishment of G protein signaling via direct Arrestin binding. We tested this hypothesis by performing Gq BRET studies in HEK cells in which βArr1 and βArr2 were deleted (Alvarez-Curto et al., 2016) and found that the effect of SBI-553 was, in fact, only modestly enhanced (Figure 2A, Tables S2/S3). We further confirmed lack of Arrestin-mediated desensitization in the above knockout cells in orthogonal assays by testing SBI-553 effects on Gi1 activation and Gq protein recruitment (Figures 1G, 1J and 2A) using the MiniGsq and MiniGsi1 G protein variants (Wan et al., 2018) and on Gq activity via standard PI hydrolysis (Figures 1H and 1L). Taken together, SBI-553 exhibits, to our knowledge, the most complex array of allosteric effects on orthosteric signaling specific to the co-bound transducer.

### NTSR1 dynamics provide insights into SBI-553 actions

The rNTSR1 is known to have very low basal activity, so to test the dynamic nature of the allosteric site and to better understand how SBI-553 might be interacting at the allosteric site, we mutated the rNTSR1 to make it constitutively active using the mutation F358A_7.42_ which has been shown to increase its basal activity (Barroso et al., 2002). Constitutively active receptor mutants spontaneously adopt conformation states that are able to recruit and activate G protein either by releasing inactive state constraints and/or forming new interactions that stabilize a more active state (Samama et al., 1993) (Roth, 2016). We surmised that if SBI-553 acts in part to stabilize the allosteric site we would potentially see an increase in Arrestin recruitment (increase in agonist activity) but also a decrease in G protein recruitment (increase antagonist activity). However, if SBI-553 interacts passively (i.e. allosteric site must be stabilized prior to SBI-553 interaction) with the constitutively active NTSR1 then the output from the SBI-553 dose response curves could putatively show little or no effect. Indeed, what we observed from the dose response curves is a potentiation of Arrestin recruitment above basal and a diminution in G protein recruitment indicating SBI-553 can in part stabilize the allosteric site (Figure 2B, Table S2).

GPCR phosphorylation can act as a regulatory mechanism in which phosphorylation rapidly initiates impairment of receptor signaling and desensitization. Many rhodopsin family GPCRs require phosphorylation by GRK(s) prior to β-Arrestin binding and subsequent β-Arrestin-directed trafficking to clathrin-coated pits and endocytic vesicles, termed phosphorylation-dependent arrestin recruitment (Nobles et al., 2011). The rNTSR1 is phosphorylated via GRK2 and GRK5 at serine and threonine residues of its intracellular loop 3 (ICL3) and C-terminus (Inagaki et al., 2015). Thus, it is conceivable that SBI-553 G protein bias could be the result of the stabilization of NTSR1 in a conformation that undergoes continuous phosphorylation precluding further recruitment of G proteins. We investigated the role of phosphorylation by mutating all identified ICL3 and C-terminus serine and threonine residues of rNTSR1 to alanine and observed minimal effects of these mutations on SBI-553 allosteric actions on Arrestin and G protein recruitment (Figure 2C, Table S2). This indicates that while phosphorylation of rNTSR1 normally occurs upon receptor activation it might only play a minor role in connection with SBI-553 allosteric effects, agonism with respect to Arrestin recruitment and antagonism with respect to G protein signaling.

### The SBI-553 allosteric binding site is located in the intracellular cavity

To determine the molecular mechanism(s) responsible for SBI-553’s unusual mode of allosterism - biased PAM antagonism, we used cryoEM to elucidate the SBI-553 allosteric site location on NTSR1. We determined the structure of rNTSR1 in complex with heterotrimeric miniGo, NTS_8-13_ with and without SBI-553 to 2.88Å and 2.89Å, respectively (Table S1). The allosteric site for SBI-553 on rNTSR1 is unambiguously located in the intracellular cavity at the interface with the G protein C-terminal helix (C*α*T) (Figure 1A and 4A).

**Figure 4.**
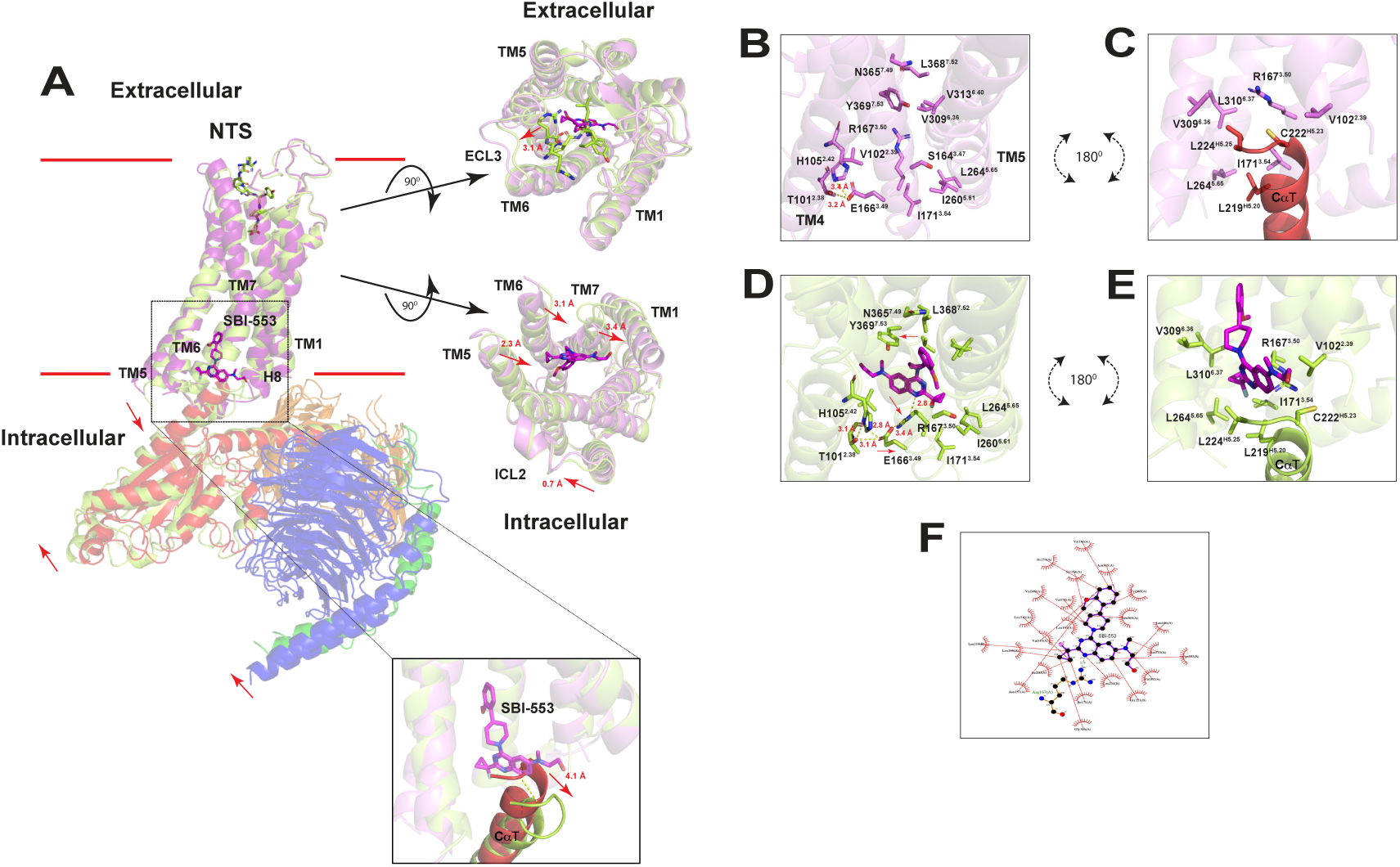
Overview of rNTSR1 architecture in the presence and absence of SBI-553 reveal insights into G protein:SBI-553 mutual permissiveness. A) Superposition of bound (light green) and unbound (light pink) NTSR1 structures reveal overall 0.79 Å RMSD (root mean squared deviation) for all Cα atoms of NTSR1. SBI-553 allosteric site is located in the intracellular cavity with its pendant phenyl ring interjecting deepest into the cavity and its quinazoline heterocyclic aromatic moiety exposed to the intracellular space. At the extracellular of NTSR1 is NTS_8-13_ in stick figures. Red line indicates approximate delineation between the transmembrane region and the extracellular and intracellular segments of rNTSR1. **Extracellular view** - looking down the receptor from the extracellular view depicting the orthosteric site to allosteric binding site. In stick figures are NTS_8-13_ (light green), SBI-553 is colored in violet. Only minor movement of TMs with the greater change seen in ECL3 which connects TM6 and TM7, distance differences between the two receptors in red lettering. **Intracellular view** - Intracellular comparison of the two structures with arrows indicating movement of TMs and loops with distance differences between the two receptors in red lettering. B) and C) Side views of NTSR1 focused on the allosteric site showing residues in the SBI-553 unbound complex with Go C- terminal helix in the intracellular cavity. D) and E) Side views of NTSR1 focused on the allosteric site showing residues in the SBI-553 bound complex with Go C- terminal helix in the intracellular cavity. NTSR1:Go:SBI-553 structure reveals remodeling of NTSR1:Go protein interface. Allosteric site residues identified undergo remodeling in the presence of SBI-553. Residues labeled with Ballesteros-Weinstein designation. F) Stick figure representation of SBI-553 with dashed lines indicating hydrophobic and charged interactions with the identified NTSR1 residues. Stick representation of SBI-553 created using Ligplot+ with the structure of NTSR1:Go:SBI-553 bound structure. Alignment was accomplished with align command using PyMOL.

Superposition of the NTSR1 unbound and bound SBI-553 NTSR1:Go complexes reveal an overall 0.79 Å RMSD (root mean squared deviation) for all Cα atoms of NTSR1 (Figure 4A). A comparison of the extracellular regions between the NTSR1:Go:SBI-553 and NTSR1:Go structures reveal this region to be relativity unchanged with very minor differences in the TMs but with some changes seen in the extracellular loops (Figure 4A). Structural changes can readily be seen at the G protein interface between NTSR1:Go:SBI-553 and NTSR1:Go structures. Significant structural changes are observed in the intracellular region of NTSR1 in the NTSR1:Go:SBI-553 complex in which TM6 and TM7 are shifted by 2.7 Å and 2.2 Å, respectively, when compared to NTSR1 in the NTSR1:Go unbound SBI-553 complex. Importantly, the effect of SBI-553 binding causes the orientation of the G protein heterotrimer to rotate/shift by ∼ 5^0^ or ∼ 3 Å relative to the unbound SBI-553 G protein heterotrimer. Despite the shift of the heterotrimeric G protein, the total interface area between the receptors and G protein remain relatively the same between the bound and unbound SBI-553 structures, 1373 Å and 1408 Å, respectively.

Allosteric sites can be transient and only available for binding of an allosteric modulator in a given receptor state as exemplified by the sodium ion binding site which is lost upon receptor activation (Katritch et al., 2014). The heretofore identified allosteric binding sites for non-sodium allosteric modulators include the extracellular vestibule, on the exterior of the 7TM bundle, and on the intracellular surface (Thal et al., 2018). For example, the M2 muscarinic acetylcholine receptor (M2R) allosteric modulator binds directly to the orthosteric site, while other Class A GPCR allosteric modulators bind to the outside of the 7TM bundle such as the P2Y Purinoceptor 1 (P2Y_1_)(Kruse et al., 2013; Zhang et al., 2015). Indeed, the SBI-553 allosteric site on NTSR1 is in a similar region shared by several previously identified allosteric modulators (Zhuang et al., 2021) (Chen et al., 2022), although SBI-553 inserts ∼ 7 Å deeper into the intracellular cavity when compared to the other allosteric modulators. This occurs principally through the piperidine and phenyl moiety extensions off the quinazoline heterocyclic aromatic base with SBI-553’s pendant phenyl ring situated furthest into the NTSR1 intracellular cavity and its quinazoline moiety exposed to the intracellular matrix and transducer interface. SBI-553 occupies ∼435 Å^2^ of buried surface area (BSA) which is on the upper end of BSA as most allosteric modulators have on average approximately 330 Å^2^ of BSA (Cheng et al., 2017; Kruse et al., 2013; Liu et al., 2017; Lu et al., 2017; Robertson et al., 2018; Zheng et al., 2016) ranging from 159 Å^2^ for AZ8838 bound to the Protease-activated receptor 2 (PAR2) (Cheng et al., 2017) to 487 Å^2^ for CMPD-15A bound to the β2 adrenergic receptor (β_2_AR) (Liu et al., 2017). SBI-553 interacts with 17 NTSR1 residues and an additional 4 residues of the Go C*α*T helix predominantly through backbone hydrophobic and sidechain aliphatic interactions (Figures 4D, 4E and 4F, Table S3).

Briefly, the methoxy substituent of the pendant phenyl ring is engaged in hydrophobic interactions with the aliphatic sidechains of L163^3.46^ and S164^3.47^. The pendant phenyl ring is engaged in hydrophobic interactions with the aliphatic sidechain of L163^3.46^ and V313^6.40^ being essentially sandwiched between these two residues. The aliphatic sidechains of residues L109^2.46^, V160^3.43^, I253^5.54^, V313^6.40^ and N365^7.49^ make hydrophobic interactions with the pendant phenyl ring forming a hydrophobic lid. Combined, the pendant phenyl ring along with its methoxy substituent essentially pin this moiety in its location. SBI-553’s piperidine moiety can be seen as having adopted a twisted chair conformation forming hydrophobic interactions with the aliphatic sidechains of V309^6.36^, Y369^7.53^, and V372^7.56^. The quinazoline heterocyclic aromatic amines of SBI-553 are sp^2^ hybridized, one of which forms a weak hydrogen bond with the amide of R167^3.50^ sidechain in an almost *π*-cation interaction. The fluorine of the cyclopropyl ring substituent off the quinazoline is also engaged in a charge interaction with the R167^3.50^ sidechain amine. Consistent with previous data, the flexibility of the piperidine ring and the addition of the cyclopropyl fluorine are contributing factors for the increased allosteric effect of SBI-553 on NTSR1, side-by-side comparison of SBI-553 and the parent compound (ML314) reveal an approximate 10-fold higher concentration was required to achieve a similar allosteric effect as seen with SBI-553 (Figure 2D, Table S2) (Ref – Pinkerton et al.). The intracellular face of SBI-553’s quinazoline heterocyclic aromatic moiety along with its fluorinated cyclopropyl and N-methylethanolamine engage Go C-terminal residues C222^H5.23^-L224^H5.25^ through a combination of charged and hydrophobic interactions with the Go C*α*T helix sidechains and the backbone.

### SBI-553 remodels the NTSR1 G protein interface

In Class A GPCRs, general structural changes associated with active state transitions include the shrinking of the extracellular vestibule, a large outward movement of TM6, an inward movement of TM5 and TM7, and a rotation of TM3 (Zhou et al., 2019). These structural changes are facilitated by “microswitches” such as the sodium ion-binding pocket, the CWxP, PIF, NPxxY, and the E/DRY motifs found within the 7TM bundle of the GPCR and link the orthosteric and transducer binding sites (Wacker et al 2012). Visual inspection of the bound and unbound SBI-553 complexes reveal 12 of the 17 NTSR1 identified residues that interact with SBI-553 have undergone either reorientation or repositioning from their unbound states (Figures 4B, 4C, 4D and 4E, Table S3). Of special note are NTSR1 residues within the E/DRY and NPxxY microswitches along with the displacement of the Go C*α*T helix from its unbound SBI-553 position within the intracellular cavity. In the unbound SBI-553 complex structure, E166^3.49^ of the E/DRY motif forms an ionic interaction with T101^2.38^ and H105^2.42^ while R167^3.50^ of the E/DRY motif is engaged in side-orientated *π*-cation interactions with Y369^7.53^ of the NPxxY motif along with charged and hydrophobic interactions with Go C*α*T helix residues C222^H5.23^ - L224^H5.25^ sidechains and backbone. Aside from Y369^7.53^, other members of the NPxxY motif are engaged in charged and hydrophobic interactions principally with members of TM6 presumably to stabilize the complex.

Noticeably, in the bound SBI-553 complex structure there is a displacement of the Go C*α*T helix when compared to the unbound SBI-553 Go complex. In addition, in the bound SBI-553 complex all the forementioned interactions are disrupted with the miniGo adopting a shifted orientation, discussed previously. NTSR1 residue E166^3.49^ sidechain in the SBI-553 bound complex is still engaged in a charged interaction with T101^2.38^ sidechain but has disengaged from H105^2.42^ forming a charged interaction with R167^3.50^ sidechain - albeit the EM density for E166^3.49^ carboxylate and hydroxyl is weak. Residue R167^3.50^ sidechain in the SBI-553 bound complex is now distorted and “pushed” to the bottom of the intracellular cavity interacting with SBI-553’s quinazoline heterocyclic aromatic moiety. Residue Y369^7.53^ sidechain has “swung” approximately 90^0^ and forms a charged interaction between its hydroxyl and the backbone amide and carbonyl of H105^2.42^ and L106^2.43^, respectively. Consistently the C*α*’s of the NPxxY motif residues are displaced by at least 2 Å from the unbound SBI-553 structure. Thus, remodeling of the NTSR1:G protein interface along with the repositioning of the G protein appears to contribute to mitigating SBI-553’s effects on Gi/o specific signal transduction.

In the BRET2 assay, SBI-553’s effect on NTSR1:Gi/o signaling was minimal in contrast to Gq/G11 signaling (Figures 1E/F/I/J, 3A and 3B). To understand the effects on Gq/G11 signaling we determined the cryoEM structure of NTSR1 in complex with our previously described Gq protein (Kim et al, Cell 2020; Cao et al Nature 2022). Despite processing NTSR1 in the identical manner as the Go complex, our attempts at determining the structure in the presence of SBI-553 yielded a complex devoid of any detectable density for SBI-553. Superposition of NTSR1 in the Gq complex with that of NTSR1 in the unbound SBI-553 Go complex revealed they superimpose well with an RMSD of 0.69 Å, with the E/DRY and NPxxY motif residues of the respective complexes are readily superimposable with only minor modelling deviations from each other. Indeed, the carbon backbone of each G protein C*α*T helix aligns well but begins to diverge upon exiting the intracellular cavity leading to an obvious deviation of C*α*T helix that further affects the superposition of the heterotrimeric complex (Figure 5A). Superposition of the Gq complex with the Go:SBI-553 complex reveals that in the Gq complex, similar to the unbound SBI-553 Go complex, the C*α*T helix would clash with SBI-553 and occlude it from engaging NTSR1(Figure 5B). However, given the lack of EM density for SBI-553, the NTSR1:Gq interface does not appear to undergo remodeling in the presence of SBI-553 and is likely excluded from the allosteric site when Gq is bound to NTSR1. Rather, SBI-553 binds to the allosteric site in the absence of Gq and then itself occludes Gq from binding to the intracellular cavity in a manner that modulates Gq signal transduction to a greater extend then that seen for Go signal transduction.

**Figure 5.**
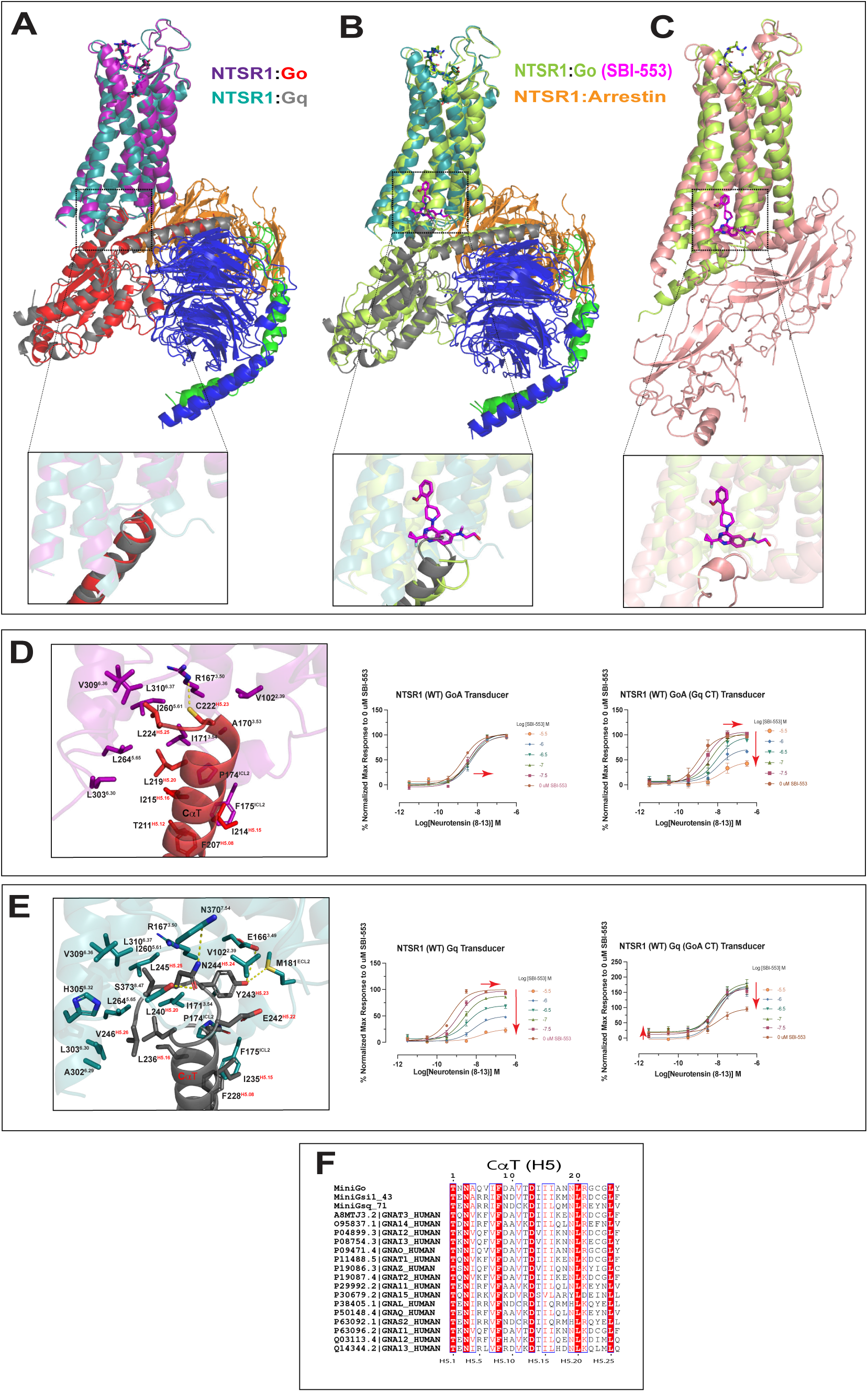
Structural alignment and functional assay reveals role of G protein C-terminus on SBI-553 permissiveness and signal modulation. A) Superposition of unbound SBI-553 structure of NTSR1:Go and NTSR1:Gq complexes. B) Superposition of bound SBI-553 structure of NTSR1:Go and NTSR1:Gq complexes. C) Superposition of bound SBI-553 structure of NTSR1:Go and NTSR1:Arrestin complexes. Go omitted for clarity. Insert are zoomed views of allosteric site with potential clash with SBI-553. D) Zoomed view of allosteric in the unbound NTSR1:Go complex, Stick figures of Go C-terminus and NTSR1 interactions. Graphs are BRET transducer assay with swapped C-terminus described in main text. E) Zoomed view of allosteric in the unbound NTSR1:Gq complex, Stick figures of Gq C-terminus and NTSR1 interactions. Graphs are BRET transducer assay with swapped C-terminus described in main text. Alignment was accomplished with align command using PyMOL. F) Sequence of alignment of G protein C*α*T as described in main text. Figure adapted from (Flock et al., 2015)using Espript server(Robert and Gouet, 2014) Given is UniProtKB name and accession number. Red arrows indicate relative effect on fitted dose-response curves with the addition of SBI-553.

Analysis of the NTSR1:G protein interfaces provides insight into SBI-553 G protein subtype selectivity. As stated earlier, the overall BSA between NTSR1 and Go protein in the bound and unbound SBI-553 complexes remained relatively unchanged despite obvious structural changes. In the unbound SBI-553 NTSR1:Go complex, NTSR1 contributed 15 residue sidechains and 593 Å^2^ BSA to the interface while the C*α*T helix contributed 10 residue sidechains and 642 Å^2^ BSA (Figure 4B and 4C). In the SBI-553 bound complex, NTSR1 contributed 17 residue sidechains and 553 Å^2^ to the shared interface while C*α*T helix contributed 8 residues sidechains and 580 Å^2^ to the shared interface (Figure 4D and 4E, Table S3). Analysis of the NTSR1:Gq complex interface revealed NTSR1 contributed 25 residue sidechains and 802 Å^2^ in BSA while the C*α*T helix contributed 12 residue sidechains and 870 Å^2^ in BSA, indicating a much broader and ‘tighter’ interface in the NTSR1:Gq complex.

The unbound SBI-553 NTSR1:Go and NTSR1:Gq structures share many conserved interactions between the respective C*α*T helix and NTSR1 transmembrane helices TM4, TM5 and TM6. Of note, are the conserved C*α*T helix residues L^H5.25^, L^H5.20^ and I/L^H5.16^ which through hydrophobic interactions use their aliphatic sidechain to interact with aliphatic sidechains of 8 NTSR1 residues (R167^3.50^, I171^3.54^, P174^ICL2^, I260^5.61^, L264^5.65^, L303^6.30^, V309^6.36^ and L310^6.37^). These NTSR1 residues are spatially in proximity with each other and form a hydrophobic ‘groove’ by which the conserved C*α*T helix residues reside (Figures 5D and 5E). In the bound SBI-553 structure, SBI-553’s fluorinated cyclopropyl moiety extends into the TM4/5/6 ‘groove’ essentially competing for the NTSR1 residues and displacing the C*α*T helix residues. However, the fluorinated cyclopropyl does not interact as extensively as the C*α*T helix residues, instead interacting with only 5 NTSR1 residues (R167^3.50^, I171^3.54^, I260^5.61^, L264^5.65^, L310^6.37^). Additionally, in the Go:SBI-553 complex, Go C*α*T residues L219^H5.20^ and L224^H5.25^ are the only residues that participate in interactions with hydrophobic “groove” sidechains interacting with only 2 NTSR1 residues, L264^5.65^ and L303^6.30^. Thus, while SBI-553 competes with Go for interactions with the hydrophobic ‘groove’ residues, it does not do so completely giving rise to the structure were both SBI-553 and Go occupy the intracellular cavity and are mutually permissive with each other.

Non-conserved residues of G protein C*α*T helix within the intracellular cavity affect SBI-553’s ability to modulate signal transduction. Noticeably absent in our Go:NTSR1 structures is the non-conserved C*α*T helix pendant Y225^H5.26^ which has been removed due to lack of convincing density in both unbound and bound SBI-553 NTSR1:Go structures. Of the other non-conserved residues within the intracellular cavity (G223^H5.24^, C222^H5.23^ and G221^H5.22^), only C222^H5.23^ makes a charged interaction with NTSR1 thorough R167^3.50^ sidechain, there are no apparent interactions with G221^H5.22^ and G223^H5.24^. Conversely, in the Gq:NTSR1 structure, non-conserved residues E242^H5.22^, Y243^H5.23^, N244^H5.24^ and V246^H5.26^ are involved in both hydrophobic and charges interactions with 8 NTSR1 residues (V102^2.39^, E166^3.49^, A170^3.53^, M181^ICL2^, A302^6.29^, H305^6.32^, N370^7.54^ and S373^8.47^) (Figure 5E). Whereas Go C*α*T intracellular cavity interactions were primarily focused to the TM4/5/6 ‘groove’, Gq’s C*α*T intracellular cavity interactions are dispersed across the intracellular cavity providing multi-point interactions that are not limited to one area of the interface.

It is widely accepted that the G*α* C*α*T plays a critical role in G protein selectivity for a given receptor (Jelinek et al., 2021) (Flock et al., 2017). Thus, it seemed reasonable that swapping the G protein C*α*T should affect SBI-553 modulation of G protein signal transduction. To test this, we swapped G protein C*α*T helix residues (D341^H5.13^ – Y354^H5.26^, D346^H5.13^ – V359^H5.26^, Go and Gq, respectively) and analyzed these swapped C*α*T G proteins via BRET2 assay. The swapping of the C*α*T helix residues did indeed affect the signal transduction such that the swapped C*α*T G proteins now produce signaling profiles similar to the unmodified G proteins, (i.e. modified Go behaves similar to unmodified Gq and modified Gq behaves similar to the unmodified Go), indicating that the C*α*T is sufficient to affect the signal transduction modulated by SBI-553. The lack of density in the Gq:NTSR1 complex structure for SBI-553, the extensive and more dispersed Gq:NTSR1 intracellular cavity interface, and convincing BRET2 data for ‘swapping’ of C*α*T residues indicate that SBI-553 and Gq are mutually exclusive to one another for occupation in the NTSR1 intracellular cavity.

Superposition of NTSR1:SBI-553 and NTSR1:Arrestin structures indicate SBI-553 and Arrestin are predicted to be mutually permissive with each other for the allosteric site. As discussed previously, the SBI-553 lead compound was identified through HCS Arrestin translocation assay and in our Arrestin recruitment assays, SBI-553 recruits Arrestin to NTSR1 in the absence of NTS (Figure 1E and 1I). Given our current structures it seemed plausible that Arrestin and SBI-553 are mutually permissive for each other in the allosteric site and/or the intracellular cavity. To address this question we superimposed the NTSR1:Arrestin structure (PDB ID 6UP7) with our NTSR1:Go:SBI-553 structure, having an 0.99 Å RMSD (Figure 5C). It can be seen the Arrestin finger loop does not inject as deeply into the NTSR1 intracellular cavity as seen with the C*α*T helix in the unbound Go and Gq structures and would not putatively clash with SBI-553. Additionally, the Arrestin finger loop would also not putatively preclude SBI- 553 from interacting with residues of the TM4/5/6 hydrophobic ‘groove’ as seen in the NTSR1:Go:SBI-553 structure such that the two (SBI-553 and Arrestin finger loop) are putatively mutually permissive with one another within the NTSR1 intracellular cavity.

## Discussion

Here, we determined the cryoEM structures of NTS8-13 bound rNTSR1 in complex with heterotrimeric miniGo, miniGq, with and without SBI-553 and unambiguously identify the allosteric site as being located in an intracellular cavity. We compared the complex of rNTSR1 with the biased allosteric modulator, SBI-553, to structures of rNTSR1 in different signaling states while also investigating SBI-553’s effect on NTSR1 signaling pathways to establish a mechanism by which NTSR1 is modulated. We provide pharmacological and structural information that illuminate an unanticipated complex mode of GPCR allosteric modulation. Pharmacological analysis revealed SBI-553 possesses a broad spectrum of allosteric modulation at NTSR1 and is a biased negative allosteric modulator with activity at Gq > G15 > Gi >> G12 and PAM-agonist activity for Arrestin recruitment. Structurally we revealed that Go G protein and SBI-553 can be mutually permissive with each other in the intracellular cavity while Gq G protein and SBI-553 are mutually exclusive with each other in the intracellular cavity. We also revealed that this intracellular cavity permissiveness is determined by the G protein C-terminus. Our data are consistent with a model in which SBI-553 is a transducer-specific biased allosteric modulator that enhances neurotensin affinity and stabilizes a state in which Arrestin translocation is potentiated while Gq and G15 signaling is inhibited and Gi1 and G12 signaling remains relatively unchanged.

We used cryoEM to determine the molecular mechanism(s) responsible for SBI-553 allosterism on NTSR1 --biased PAM antagonism. Initially we pursued x-ray crystallography. However, this approach did not yield the location of the allosteric site as many crystals either failed to diffract or there was no convincing extra density that could be attributed to SBI-553. Hence, we turned to cryoEM to elucidate the SBI-553 allosteric site location on NTSR1. Attempts to identify the allosteric site using an NTSR1:Arrestin complex was unsuccessful due to low resolution. We then turned to structure determination using a NTSR1:G protein complex. To do this, we screened via BRET assay the available NTSR1 transducerome (Olsen et al., 2020) to identify a complex which would be amenable for structure determination, i.e. results that showed a degree of agonism in the presence of SBI-553 but in the absence of NTS resulting in complex structures of rNTSR1 in complex with heterotrimeric miniGo, miniGq, NTS8-13 with and without SBI-553.

The simplest model for allosterism is one of conformational selection between two states, active and inactive. In the allosteric two state model, the ligand free receptor exists in both inactive and active states (R and R*) with two ligands, A (orthosteric) and B (allosteric), binding to different sites on the receptor with the allosteric modulator mediating the change between inactive and active states (Ref - Hall, 2000). High-resolution GPCR structures and other studies have provided snapshots of GPCR dynamics noting that a two-state model explanation is unlikely to accommodate a receptor’s full conformational repertoire. Thus, it is widely accepted that GPCRs are highly dynamic and can adopt multiple active and inactive states leading to distinct outcomes. These states can be differentially stabilized by not only orthosteric ligands (agonists, antagonists) but also by allosteric ligands that bind at spatially distinct sites from the orthosteric site (PAM, NAM Ago-PAM, etc.) (Canals et al., 2011). To date, small molecule allosteric modulators of class A GPCRs have been shown to bind in three separate regions of GPCRs including the extracellular vestibule, outside of the 7TM bundle (in multiple locations), and to the intracellular surface. A common theme, as might be expected, is that PAMs promote the stabilization or transition of the active state while NAMs promote the inactive state through mechanism that prevent the transition to the active state or by occluding the binding of transducers. For example, in the M2R, binding of the orthosteric agonist iperoxo promotes closure of the extracellular vestibule, and binding of the PAM LY2119620 further contracts the orthosteric binding site. Conversely, NAMs that would bind to the extracellular vestibule would be expected to prevent its closure. Allosteric modulators that bind to the outside of the 7TM bundle either act by stabilizing (PAM) or preventing the rotation of TM3 (NAM). While modulators that bind to the intracellular surface prevent the binding of transducers can in some instances actually stabilize the active state. While there exist structures of class A GPCRs in complex with PAM or NAM allosteric modulators, in most cases, there is still a lack of understanding as to how and as to what extent these modulators affect other signaling pathway due to lack of rigorous pharmacological characterization so none of these allosteric modulators are known to be biased allosteric modulators which preference the receptor to a given pathway.

The therapeutic utility of biased agonists and allosteric modulators for GPCRs is an emerging area of GPCR drug design and discovery. Key pharmacologic characteristics of GPCR allosterism include 1) improved selectivity due to either greater sequence divergence between receptor subtypes and/or subtype-selective cooperativity, 2) probe dependence in which the magnitude and direction of the allosteric effect can change depending on the ligand, and 3) the potential for biasing the receptor towards or away from a given pathway. There are known examples of G protein biased signaling allosteric modulators of a class A GPCRs such as BMS 986187 for the delta opioid (DOR) but there are fewer examples of Arrestin biased allosteric modulators. One such example is the use of pyrimidinyl biphenylureas that act as PAMs to promote binding of the agonist CP55940 to the Cannabinoid Receptor 1 (CB1) stimulating receptor internalization and Arrestin recruitment in a biased manner. Frequently allosteric modulators have low affinity and/or complex interactions with the receptor contributing to the lack of understanding as to their structural basis. However, despite not having complex structures, there is still good rationale of the clinical relevance and potential for Arrestin biased drugs. For example, carvedilol is a nonsubtype-selective β-adrenergic receptor (βAR) antagonist has been proven to be particularly effective in the treatment of heart failure while later identified as arrestin biased at the β2AR.

The unusual profile of the biased signaling effects of SBI-553 predict an interesting panorama of activity *in vivo*. On one hand, SBI-553 may produce direct β-Arrestin agonism concomitant with Gq inverse agonism, provided sufficient basal tone is present at the Gq pathway. On the other hand, SBI-553 will modify the quality of the efficacy of the natural agonist neurotensin producing a NTSR1 biased toward β-Arrestin stimulation and away from Gq protein stimulation. The enhancement of neurotensin affinity also will produce a unique profile of G protein antagonism as PAM antagonists enhance their antagonist potency in the presence of the natural agonist. This profile leads to uniquely active antagonism, as seen in the case of the 5-HT_3_ PAM antagonist for the treatment of nausea, and an extended duration of action *in vivo* (Kenakin and Strachan, 2018). Taken together our findings reveal a complex mode of GPCR allosteric modulation and provide a structural basis for these actions. In a recent manuscript, the therapeutic consequences of this unusual mode of allosteric modulation are revealed for psychostimulant drugs (Slosky et al., 2020). Our findings provide the structural details for biased allosteric modulation and should facilitate the design and discovery of selective biased allosteric modulators for NTSR1 and perhaps other GPCRs.

## Accession Numbers

Coordinates of NTSR1 complexes and EM maps have been deposited in the Protein Data Bank and the Electron Microscopy Data Bank under the accession codes 8FN1 (EMD-29303), 8FN0 (EMD-29302), and 8FMZ (EMD-29301) for the NTSR1 Go, Go:SBI-553 and Gq complexes, respectively.

## Acknowledgements

This work was supported by NIH Grant RO1MH112205, the NIMH Psychoactive Drug Screening Program Contract and the Michael Hooker Distinguished Chair of Pharmacology (B.L.R.). We gratefully acknowledge the UNC CryoEM Core Facility along with the UNC Flow Cytometry Core Facility both facilities are supported in part by P30 CA016086 Cancer Center Core Support Grant to the UNC Lineberger Comprehensive Cancer Center.

## Author Contributions

B.E.K. designed experiments, expressed, screened and prepared grids for cryoEM constructs, processed the cryoEM data and refined the structure, performed ligand binding and functional assays, analyzed all data, and prepared the manuscript. J.F.D and R.H.J.O. optimized and performed functional experiments including BRET1 (Recruitment) and BRET2 assays. HJ.K. optimized and performed saturation, association and dissociation experiments. S.T.S. optimized and performed IP3 functional experiments. J.F.F. prepared the cryoEM grids, collected and processed cryoEM data. R.T.S. and T.K. analyzed allosteric data and provided guidance on allosteric functional assays. A.B.P., L.M.S., L.S.B., M.G.C. provided ML314, SBI-553 and other analogs for testing in functional assays and structure determination. L.M.S., L.S.B., M.G.C. provided input for assay analysis. All authors contributed to the writing and/or editing of the manuscript. B.L.R was responsible for the overall project strategy and management.

## Methods

### NTSR1 constructs

The baculovirus construct rNTSR1 consisted of the hemagglutinin signal peptide and the Flag tag, a deca-histidine tag was placed at the N-terminus followed by a prescission protease recognition site followed by the A86L, G215A, V360A thermostabilized rat NTSR1 (T43-Y424). All function assays were performed either with WT human, Rat NTSR1 or the cryoEM construct. Assay composition is defined in the figure legend. DNA was synthesized by Integrated DNA Technologies (www.idtdna.com) and subcloned into a modified pFastBac1 vector using restriction sites of AscI and FseI.

### Construction and expression of the receptor-G protein complex

For the miniGq and miniGo heterotrimeric complexes, we used the same constructs as used in the 5-HT_2A_R-miniGq and 5-HT_5A_R-miniGo complexes. The Bac-to-Bac baculovirus expression system (Invitrogen) was used to generate the recombinant baculovirus for protein expression. Baculoviruses of corresponding miniGo-Gβ1*γ*2 or miniGq-Gβ1*γ*2 and scFv16 were co-expressed by infecting Sf9 cells at a density of 2x10^6^ cells per ml at MOI ratio of 3:1.5:1.25, respectively. Cells were harvested by centrifugation 48 h post-infection and stored at -80°C for future use.

### Receptor-G protein complex purification

The cell pellet of the rNTSR1-miniGo-scFv16 complex was thawed on ice and incubated with a buffer containing 20 mM HEPES pH 7.5, 50 mM NaCl, 10 mM MgCl_2_, 20 mM KCl, 5 mM CaCl_2_, proteinase inhibitor, 40 units apyrase, and 5 uM NTS_8-13_ at room temperature. After 1.5 h, the cell suspension was homogenized, membrane was collected by centrifugation at 100,000 x g for 35 min and solubilized using 20 mM HEPES pH 7.5, 100 mM NaCl, 10% (w/v) glycerol, 0.5% (w/v) lauryl maltose neopentyl glycol (LMNG, Anatrace, NG310), 0.05% (w/v) cholesteryl hemisuccinate (CHS), 10 mM MgCl_2_, 20 mM KCl, 5 mM CaCl_2_, proteinase inhibitor, 40 units apyrase, and 5 uM NTS_8-13_ for 5 h at 4°C. The solubilized proteins in the supernatants were isolated by ultra-centrifugation at 100,000 x g for 45 min and then incubated overnight at 4°C with TALON IMAC resin and 20 mM imidazole. The resin was collected next day and washed with 25 column volumes 20 mM HEPES pH 7.5, 100 mM NaCl, 20 mM imidazole, 0.01% (w/v) LMNG, 0.001% (w/v) CHS and 10 µM NTS_8-13_. The protein was then eluted using the same buffer supplemented with 250 mM imidazole. Eluted protein was concentrated and subjected to size-exclusion chromatography on a Superdex 200 Increase 10/300 column (GE Healthcare, 289909944) that was pre-equilibrated with 20 mM HEPES pH 7.5, 100 mM NaCl, 5 µM NTS_8-13_, 0.00075% (w/v) LMNG, 0.00025 (w/v) glyco-diosgenin (GDN, Anatrace, GDN101), and 0.00075% (w/v) CHS. Peak fractions were pooled then divided with one part supplemented with 50 uM SBI-553 and incubated for 2 h at 4 degrees then both parts were concentrated to 5 mg ml^-1^.

### CryoEM data collection, 3D reconstitution, model building and refinement

The cryoEM data collection and processing of the rNTSR1 complexes follows a published protocol (Peck et al., 2022). The samples (3.2 µl) were applied to glow-discharged Quantifoil R1.2/1.3 Au300 holey carbon grids (Ted Pella) individually and were flash-frozen in a liquid ethane/propane (40/60) mixture using a Vitrobot Mark IV (FEI) set at 4°C and 100% humidity with a blot time range from 3.0 to 4.5 s. Images were collected using a 200 keV Talos Artica with a Gatan K3 direct electron detector at a physical pixel size of 0.88 Å. Micrograph recorded movies were automatically collected using SerialEM using a multishot array (Mastronarde, 2005) Data were collected at an exposure dose rate of ∼15 electrons/pixel/second as recorded from counting mode. Images were recorded for ∼1.7-2.7 seconds in 60 subframes to give a total exposure dose of ∼45 electrons per Å^2^. Following manual inspection and curation of the micrographs, particles from each dataset were selected using the Blob particle picker and initial 2D classification yielded templates for subsequent template picking. After one round of two-dimensional classification and selection in cryoSPARC, a subset of the selected particles was used as a training set for Topaz and the particles were re-picked from the micrographs using Topaz (Bepler et al., 2020) and subjected to two-dimensional classification and three-dimensional classification. The select classified picked particle coordinates from the three sets were next merged yielding a subset of unique particles that survived 2D classification (i.e. duplicates were removed with a radius of 75 pixels). All subsequent three-dimensional classification and refinement steps were performed within cryoSPARC (Punjani et al., 2020) Multiple rounds of multi-terence refinement resolved the final stack of particles that produced a map with a resolution reported in **Table S1** (by FSC using the 0.143 Å cut-off criterion) (Rosenthal and Henderson, 2003) after Global CTF refinement and post-processing including soft masking, B-factor sharpening in cryoSPARC, and filtering by local resolution (Heymann and Belnap, 2007) to generate the post-processed sharpened map. Alternative post-sharpening was performed on the two half-maps using deepEMhancer (Sanchez-Garcia et al., 2021) For more details see Table S1 and Figure S2.

Maps from deepEMhancer were used for map building, refinement, and subsequent structural interpretation. The rNTSR1 crystal structure (PDB ID 4XES) was used as the initial model for the rNTSR1-miniGo +/- SBI-553 and rNTSR1- miniGq complexes and docked into the cryoEM map using Chimera (Pettersen et al., 2004) followed by initial rigid body and simulated annealing then iterative manual adjustment in COOT (Emsley and Cowtan, 2004) and Phenix.real_space_refine in Phenix (Adams et al., 2010) The model statistics were validated using Molprobity. Structural figures were prepared using Chimera or Pymol (https://pymol.org/2/).

Calculation of SBI-553 Buried Surface Area (BSA) was accomplished using the areaimol program of the CCP4 suite.

### Radioligand Association and Dissociation Assay

Radioligand dissociation and association assays were performed in parallel utilizing the same concentrations of radioligand, membrane preparations, and binding buffer (50 mM Tris, 0.2% (w/v) BSA, pH 7.4). All assays utilized 1 nM of radioligand ([^3^H]-Neurotensin, PerkinElmer). For dissociation assays, membranes were pre-incubated with radioligand for at least 30 min at RT before the addition of 10 μl of 10 μM excess cold neurotensin and 100 nM SBI-553 to the 200 μL membrane suspension at designated time points. For association experiments, 100 μL of radioligand (± 3 μM SBI) was added to 100 μL membrane suspensions at designated time points. At time = 0 min, plates were harvested by vacuum filtration onto 0.3% (v/v) polyethyleneimine pre-soaked 96- well filter mats (Perkin Elmer) using a 96-well Filtermate harvester, followed by three washes of cold wash buffer (50 mM Tris, pH 7.4). Scintillation (Meltilex) cocktail (Perkin Elmer) was melted onto dried filters and radioactivity was counted using a Wallac Trilux MicroBeta counter (PerkinElmer). Data were analyzed using ‘‘Dissociation – One phase exponential decay’’ or ‘‘Association kinetics –one conc. of hot’’ in Graphpad Prism 8.0 (Graphpad Software Inc., San Diego, CA).

### Arrestin and Gq Bioluminescence Resonance Energy Transfer (BRET) Recruitment Assays

To measure β-Arrestin and G protein recruitment (BRET), HEK293T cells (ATCC CRL-11268) maintained in DMEM containing 10% (v/v) dialyzed FBS, 1 IU ml^-1^ Penicillin G, and 100 ug ml^-1^ Streptomycin were passed to 10 cm dishes and co-transfected using TransIT (Mirus Bio) in an approximate 1:2.5 ratio with NTSR1 containing C-terminal *Renilla* luciferase (*R*Luc) and Venus-tagged N-terminal β-Arrestin2 or Venus-tagged N-terminal MiniG protein containing an N-terminal nuclear export signal, respectively (Wan et al., 2018). After at least 24 hours, transfected cells were plated in poly-lysine coated 96-well white clear bottom cell culture plates in plating media (DMEM containing 1% (v/v) dialyzed FBS, 1 IU ml^-1^ Penicillin G, and 100 ug ml^-1^ Streptomycin) at a density of 40,000 cells in 200 μL per well and incubated overnight.

The following day, media was aspirated, and cells were washed once with 60 μL of drug buffer (1X HBSS, 20 mM HEPES, pH 7.4). Then 60 μL of drug buffer was added per well and drug stimulation was performed with the addition of 15 μL of 6X drug dilution of SBI-553 in drug dilution buffer (1X HBSS, 20 mM HEPES, 0.3% (w/v) BSA, 0.03% (w/v) ascorbic acid, pH 7.4) per well and incubated at RT. After 80 minutes of incubation, 10 μL of the *R*Luc substrate, coelenterazine h (Promega) at 5 μM final concentration was added per well. After an additional 5 minutes, 15 μL of 6X drug dilution of NTS8-13 in drug dilution buffer was added to the each well, plates were read at 90 minutes post start of incubation for both luminescence at 485 nm and fluorescent eYFP emission at 530 nm for 1 second per well using a Mithras LB940 (Berthold Technologies). Plates were read for multiple time points up to 30 minutes. The BRET ratio of eYFP/*R*Luc was calculated per well and the net BRET ratio was calculated by subtracting the eYFP/*R*Luc ratio per well from the eYFP/*R*Luc ratio in wells without Venus-β-Arrestin present. The net BRET ratio were normalized to the no drug addition (NTS only) data and plotted as a function of drug concentration using Graphpad Prism 8 (Graphpad Software Inc., San Diego, CA).

To measure NTSR1-mediated G protein dissociation (BRET2), procedures were similar to NTSR1-mediated β-Arrestin2 recruitment, except HEK293T cells were co-transfected in a 1:1:1:1 ratio of Gα-RLuc, Gβ1, GFP_2_-Gγ2, and NTSR1, respectively (750 ng pcDNA3.1 each). G protein dissociation BRET^2^ assays utilized 10 μL of the *R*Luc substrate Coelenterazine 400a (Nanolight, 5 μM final concentration), incubated for 10 minutes, and read for luminescence at 400 nm and fluorescent GFP_2_ emission at 515 nm for 1 second per well using a Mithras LB940. The ratio of GFP_2_/RLuc was calculated per well and plotted as a function of drug concentration using Graphpad Prism 8 (Graphpad Software Inc., San Diego, CA).

### Inositol 1,4,5-trisphosphate (IP(3)) Scintillation Proximity Assay

Scintillation proximity assays of inositol phosphate production were carried out as previously described with some adaptations. In brief, HEK293T maintained in Dulbecco’s Modification of Eagle Medium (DMEM, VWR) supplemented with 10% (v/v) fetal bovine serum (FBS, VWR), 100 U mL^-1^ penicillin, 100 μg mL^-1^ streptomycin and 5% CO2 atmosphere at 37^0^ C were transiently transfected with wild type human and rat NTSR1 in pcDNA3.1 (4 μg DNA per 75% confluent 10 cm dish) via hybridization with polyethylenimine (PEI, Polysciences, Inc.). The following day, cells were seeded in 96-well poly-lysine (Sigma Aldrich) coated plates (Greiner) at a density of 80K cells in 100 μL well^-1^ of inositol-free DMEM (Caisson Labs) supplemented with 5% dialyzed FBS (VWR). 90 min after seeding, 20 μL of inositol-free DMEM containing 1 μCi [^3^H]myo-inositol (Perkin Elmer) was added to each well and incubated for an additional 18 h. Medium was aspirated and replaced with 50 μl well^-1^ of 1X Hanks balanced salt solution (HBSS), 20 mM HEPES, 24 mM NaHCO3, 11 mM glucose, 15 mM LiCl, pH 7.4 containing 0 to 10 μM SBI-553. After 1 h incubation in 5% CO2 atmosphere at 37^0^ C, 50 μL well^-1^ of 2X neurotensin 1-13 (Tocris) solutions were added to the appropriate well and incubated for an additional 30 min. Drug solutions were aspirated and replaced with 40 μL well^-1^ 50 mM ice cold formic acid then incubated at 4^0^ C for 1 h. During which, RNA-binding YSi beads (Perkin Elmer) were diluted in double-distilled water to 100 μg uL^-1^, 75 μL of beads was added to each well of a flexible 96-well scintillation plate (Perkin Elmer). After 1 hr, the formic acid solution was transferred from the assay plates to the scintillation plates, mixed, sealed, and incubated at 4^0^ C for 2 h before counting with a MicroBeta scintillation top counter (Perkin Elmer).

## SI Titles and Legends

**Figure S1.**
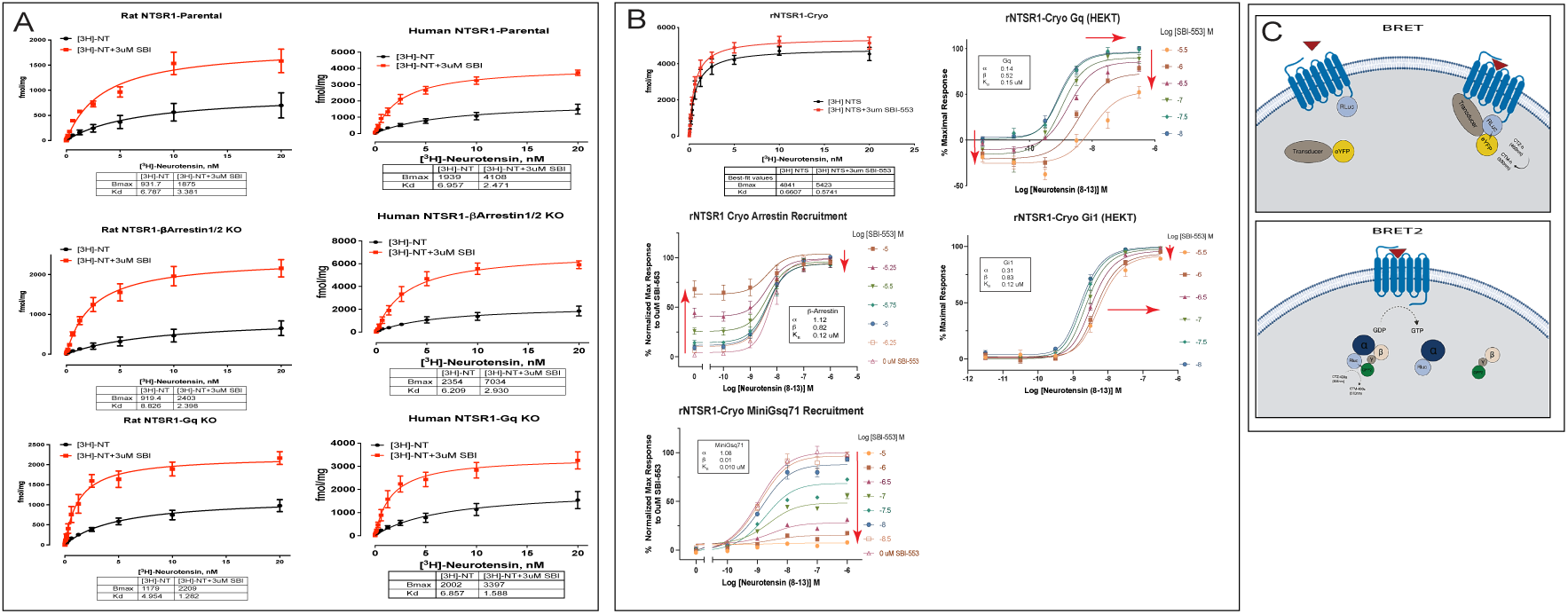
Functional characterization of rNTSR1, hNTSR1, and NTSR1 Cryo constructs. Panel A. rNTSR1 and hNTSR1 saturation binding experiments with increasing concentrations of [^3^H]NTS in the presence (red) and absence (black) of constant concentration of 3 μM SBI-553 using Parental, βarr1/2, and G11/Gq Knockout cells lines, respectively. **Panel B)** rNTSR1-Cryo construct saturation binding experiment using increasing concentrations of [^3^H]NTS in the presence (red) and absence (black) of constant concentration of 3 μM SBI-553. Graphs of dose response curves accomplished using the rNTSR1-Cryo as described in the main text. **Panel C** – Bioluminescence Resonance Energy Transfer (BRET/BRET2) Experiments. In BRET experiments, a *renilla* luciferase (*R*Luc) was placed at the C-terminus of NTSR1 while β-Arrestin2 and MiniG proteins were N-terminally tagged with “Venus” yellow fluorescent protein (YFP). NTSR1 G protein GEF activity was measured using the BRET2 G protein dissociation assay. In the BRET2 assay the *Rluc* is a chimera with the G protein α subunit (Gα) while Green Fluorescent Protein (GFP) is located on the N-terminus of the G protein γ (Gγ) subunit. Activation of NTSR1 results in a decreased net BRET signal due to nucleotide exchange and the dissociation of G protein into α and βγ subunits. Data are presented as mean values ± SEM with a minimum of two technical replicates and N = 3 biological replicates. Insert in each graph are the allosteric parameters required to fit the data using the Allosteric EC50 Shift and Black Leff Ehlert equation for PAM, General Least squares fit in Graphpad Prism 8.0 (Graphpad Software Inc., San Diego, CA). Red arrows indicate relative effect on fitted dose-response curves with the addition of SBI- 553.

**Figure S2.**
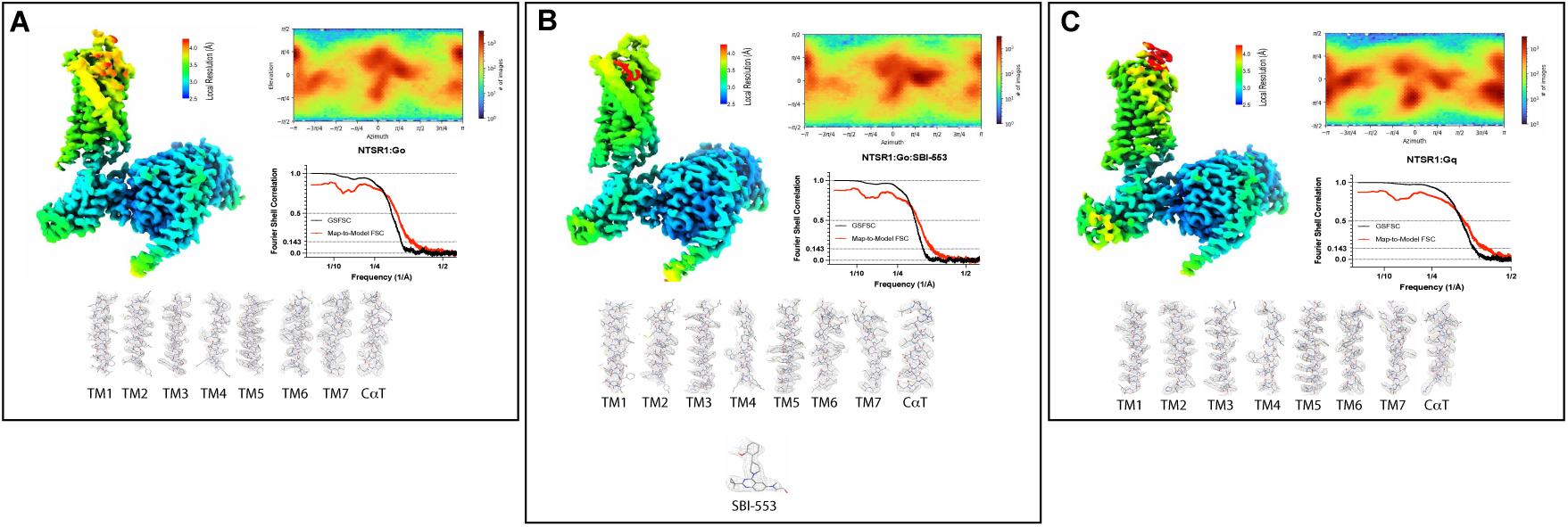
CryoEM data-processing of NTSR1 complexes. **Panel A) NTSR1:Go, B) NTSR1:Go:SBI-553, C) NTSR1:Gq - GSFSC plot of auto-**masked final map (black) and map-to-model real-space cross correlation (red) as calculated from phenix.mtriage. Direction distribution and local resolution estimation heat maps. Local cryo-EM density maps of NTSR1 TM1-7, SBI-553, and G protein helix *α*5 (C*α*T).

**Table S1.**
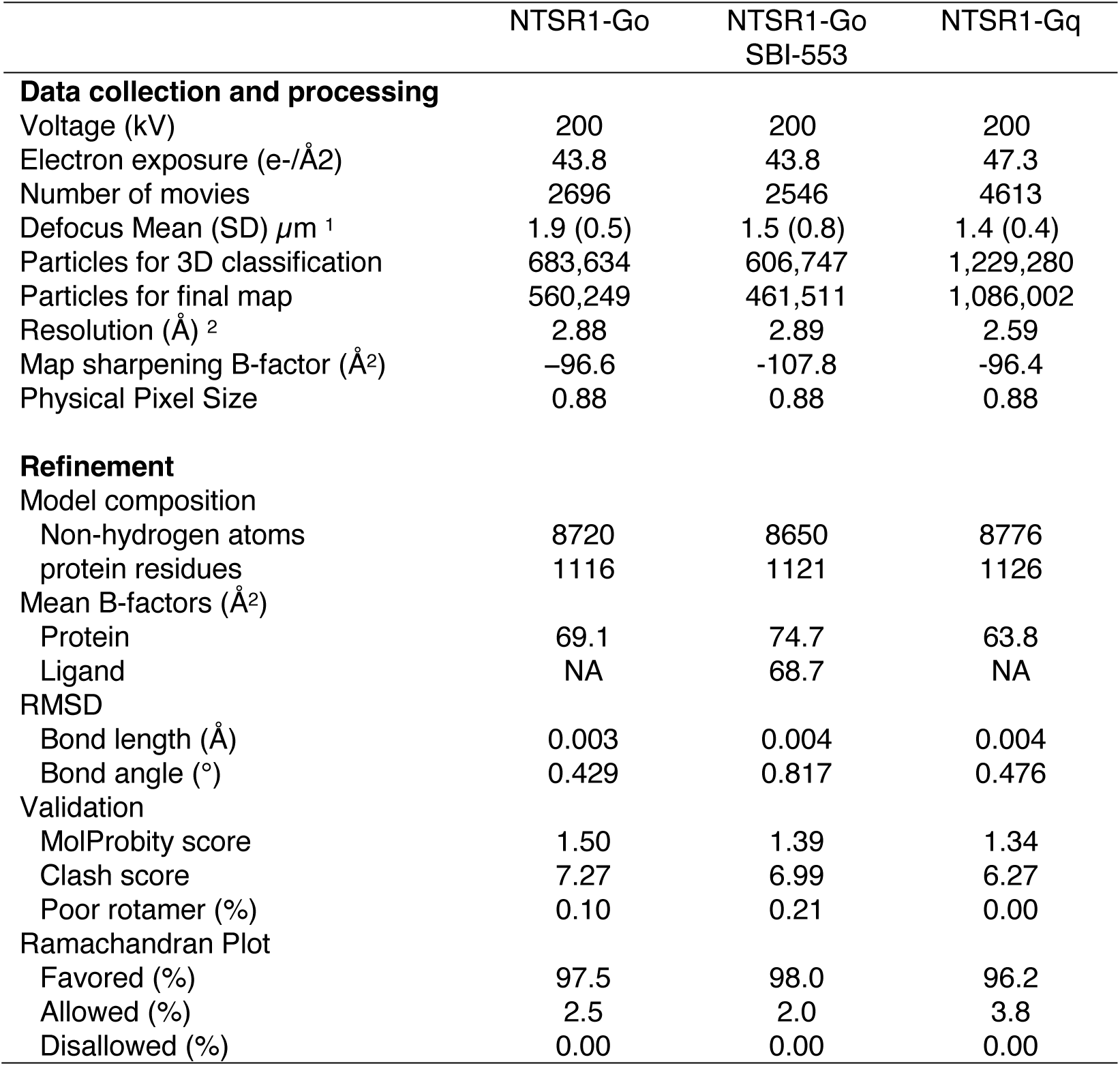
Cryo-EM data collection, refinement and validation statistics

**Table S2.**
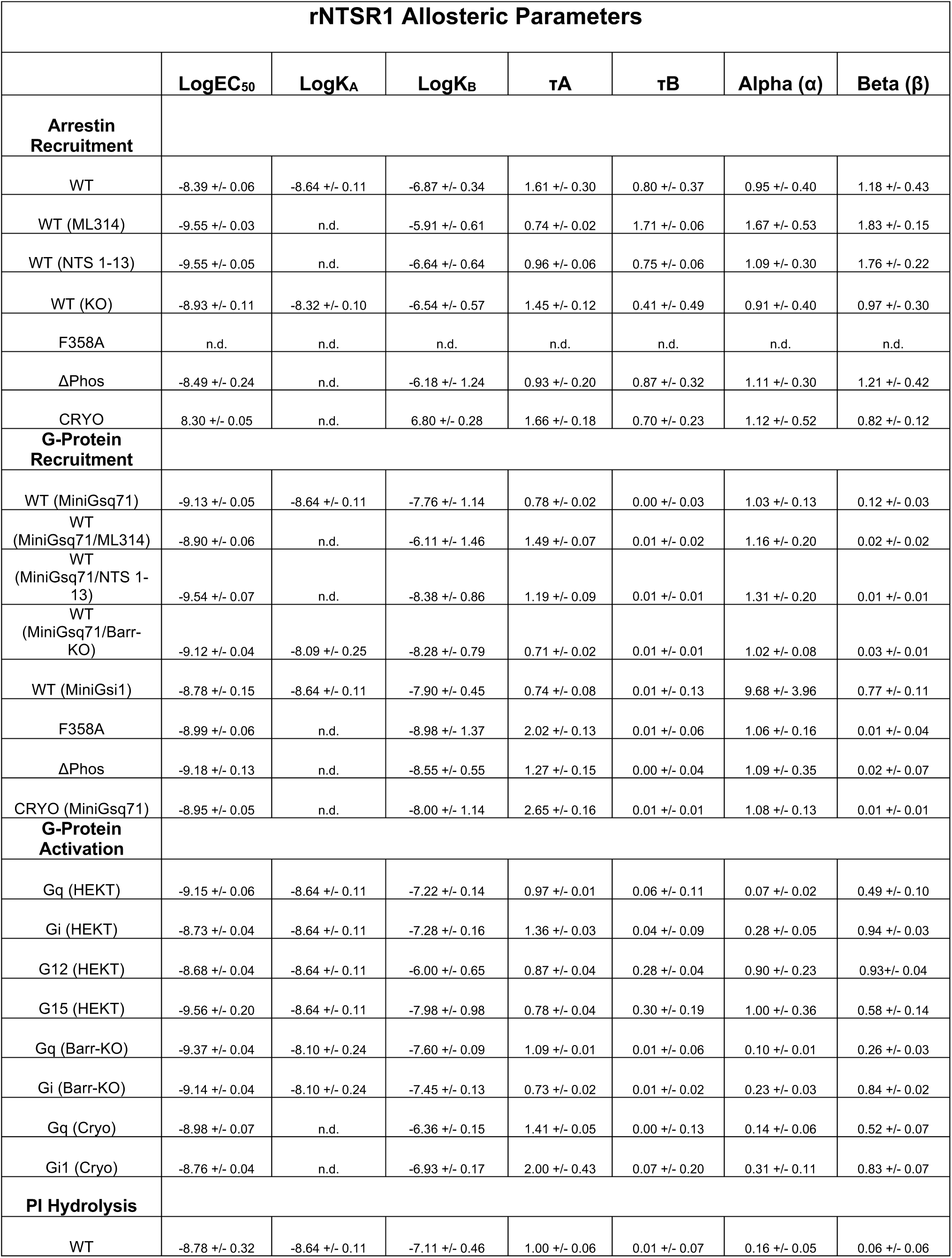
Allosteric Parameters –. Rat NTSR1 (rNTSR1) derived allosteric parameters from fitting of dose response curves of indicated assay using the Allosteric EC50 Shift and Black Leff Ehlert equation for PAM, General Least squares fit in Graphpad Prism 8.0 (Graphpad Software Inc., San Diego, CA). Data are presented as mean values ± SEM with a minimum of two technical replicates and N = 3 biological replicates.

**Table S3.**
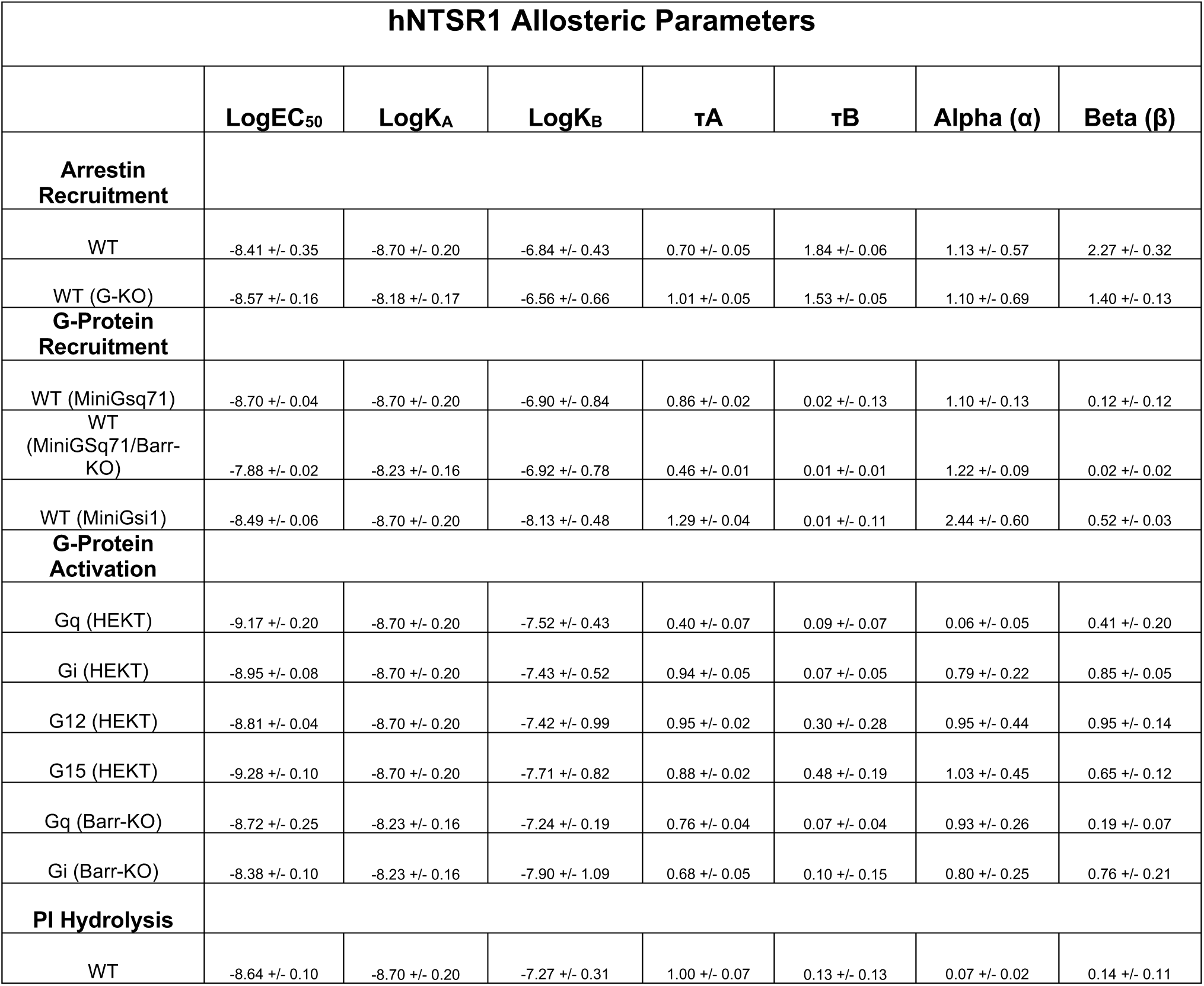
Allosteric Parameters –. Human NTSR1 (hNTSR1) derived allosteric parameters from fitting of dose response curves of indicated assay using the Allosteric EC50 Shift and Black Leff Ehlert equation for PAM, General Least squares fit in Graphpad Prism 8.0 (Graphpad Software Inc., San Diego, CA). Data are presented as mean values ± SEM with a minimum of two technical replicates and N = 3 biological replicates.

**Table S4.**
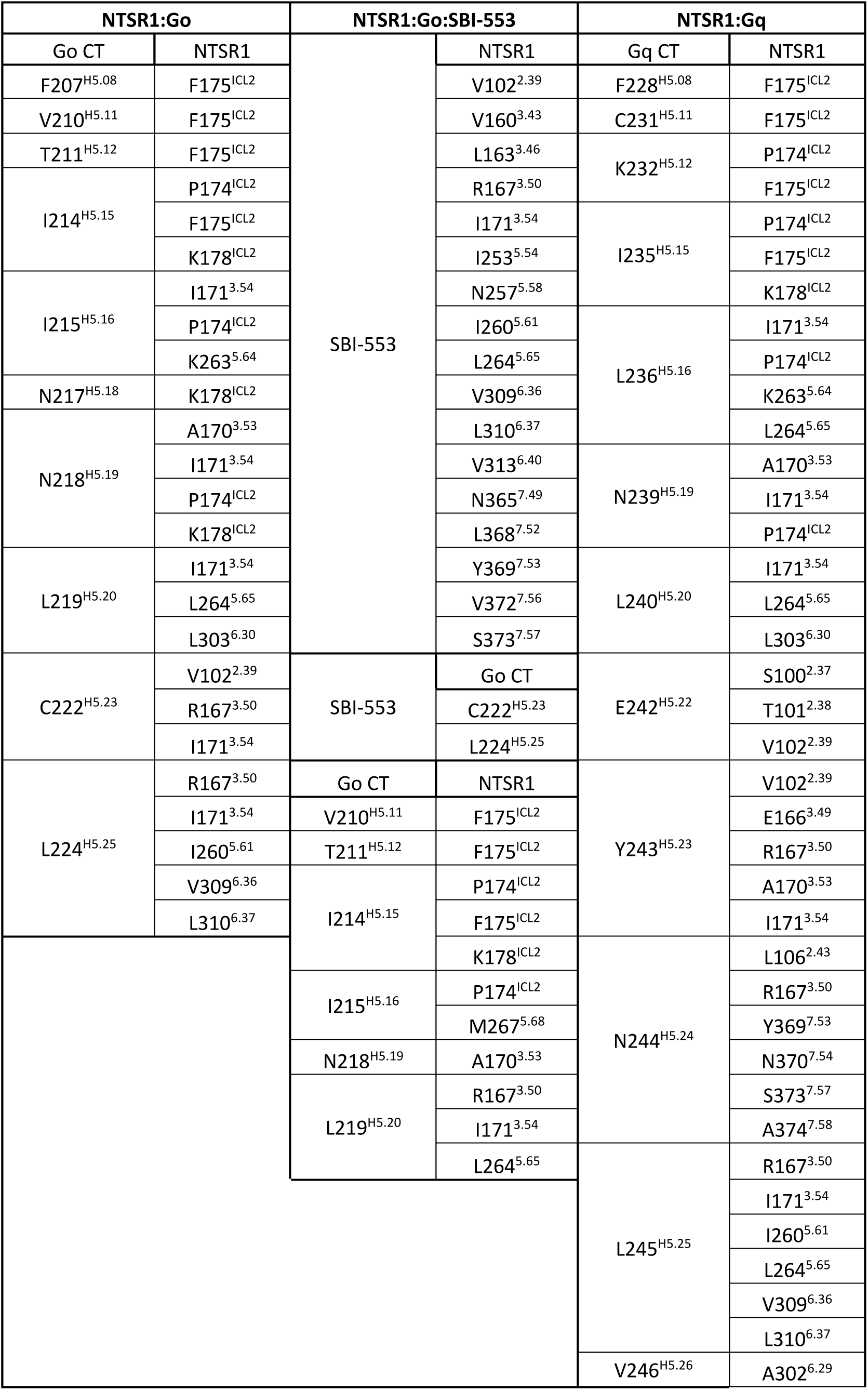
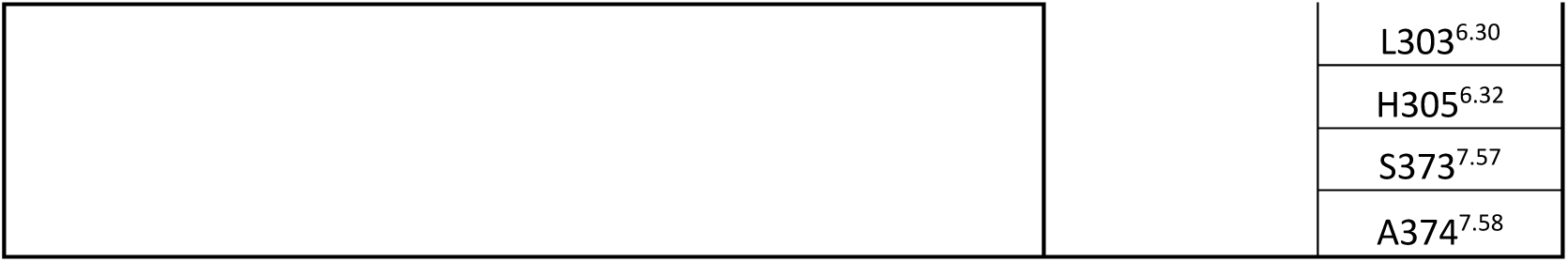
NTSR1, SBI-553 and G protein Cατ Sidechain Interactions. Sidechain interactions of identified residues with distances of less than 5 Å. Backbone interactions and sidechains with minimal density have been excluded.

